# Haplotype-resolve genome assembly and resequencing provide insights into the origin and domestication of modern rose

**DOI:** 10.1101/2023.06.02.543351

**Authors:** Zhao Zhang, Yang Liu, Tuo Yang, Shan Wu, Honghe Sun, Jie Wu, Yonghong Li, Yi Zheng, Haoran Ren, Yuyong Yang, Shaochuan Shi, Wenyan Wang, Qi Pan, Lijuan Lian, Shaowen Duan, Yingxiong Zhu, Youming Cai, Hougao Zhou, Hao Zhang, Kaixue Tang, Jiaopeng Cui, Dan Gao, Liyang Chen, Yunhe Jiang, Xiaoming Sun, Xiaofeng Zhou, Zhangjun Fei, Nan Ma, Junping Gao

## Abstract

Modern rose (*Rosa hybrida*) is a recently formed interspecific hybrid and has become one of the most important and widely cultivated ornamentals. Here, we report the haplotype-resolved chromosome-scale genome assembly of the tetraploid *R. hybrida* ‘Samantha’ and a genome variation map of 233 *Rosa* accessions involving various wild species, and old and modern cultivars. The two subgenomes of ‘Samantha’ show no significant global bias in gene loss but substantial expression bias, and homoeologous exchanges are frequently observed between subgenomes. Population genomic and genomic composition analyses reveal the contributions of wild *Rosa* species to modern roses, and highlight that *R. odorata* and its derived cultivars contribute more to modern roses than *R. chinensis* ‘Old Blush’. Furthermore, selective sweeps during modern rose breeding associated with major agronomic traits, including continuous flowering, floral organ identity, flower senescence, and disease resistance, are identified. This study provides insights into the genetic basis of modern rose origin and breeding history, and offers unprecedented genomic resources for rose improvement.

## Introduction

Rose is among the most important ornamental plants and provides a precious source of natural scent, generating billions of dollars in trade worldwide annually. It has been cultivated for thousands of years in China and Europe^1^, separately but coincidentally. The ancient rose cultivars, or so-called old roses, originated from reticulate evolution among several wild species within the genus *Rosa* of different ploidy levels^2^. As ancestors of modern rose, the wild species that have been passed down for thousands of years exhibit important agronomic/ornamental traits, including double flower, continuous flowering, unique scent, flower shape, inflorescence type, growth vigor and abiotic/biotic stress tolerance^3, 4^. These wild species exhibit a broad geographic distribution, with *R. odorata* var. *gigantea*, *R. chinensis* var. *spontanea*, *R. chinensis* ‘Old Blush’, *R. wichuraiana, R. rugosa* and *R. fedtschenkoana* predominantly found in China, *R. moschata* in East Asia, and *R. gallica* and *R. canina* throughout Europe^1^.

Since the 18^th^ century, several diploid Chinese old cultivars including *R. chinensis* ‘Slater’s Crimson China’, *R. chinensis* ‘Old Blush’, *R. odorata* ‘Hume’s Blush Tea-scented China’, and *R. odorata* ‘Park’s Yellow Tea-scented China’ had been brought to Europe, wherein they were crossed with European old cultivars and other wild accessions^3^, producing a series of intermediate types that include Bourbon, Noisette, Portland, Tea, Hybrid Perpetual etc. (Supplementary Note 1). In 1867, the hybridization effort met with great success when a hybrid named ‘La France’ was created by the rosarian Jean-Baptiste André Guillot. This hybrid rose, which combined the good traits from Chinese and European old roses, has since become recognized as the class of modern rose (referred to as *Rosa hybrida*). The modern rose shows great phenotypic variation, as well as genetic potential for yield, quality, phenological adaptability, making it quickly predominate in rose cultivation^4, 5^. Nowadays, over 35,000 modern rose cultivars, of which most are tetraploid, have been cultivated worldwide. These modern roses have a wide variety of colors, flower shapes and scent, as well as different plant architectures, all of which are important traits and need to be better understood.

According to the documentary records, at least 8-20 wild species and old cultivars with different ploidy levels were involved in the creation of modern roses^3^. The high heterozygosity resulted from the reticulate evolution, a high percentage of repetitive sequences, and the polyploidy all challenge the deciphering of the mystery of the genomic composition of modern roses. The genome of a doubled haploid (DH) line of *R. chinensis* ‘Old Blush’, which has been considered as one of the important progenitors of modern rose, has become available recently^6, 7^. However, the genomic information of tetraploid modern roses is scarce, and their origin, evolution, and improvement remain to be investigated. Here we present a haplotype-resolved high-quality genome assembly of a tetraploid modern rose cultivar ‘Samantha’ and genome resequencing of 214 rose accessions involving wild species, and old and modern rose cultivars. We present subgenome features of the modern rose, characterize the ancestral contributions of initial founder species, and identify selective sweeps during modern rose breeding history that are associated with important traits. Altogether, our work shed light on the complex ancestry of modern roses and how human selection has reshaped rose genome in a short period of less than two hundred years. Our results further provide valuable information for understanding the genetic basis of ornamental and cultivated traits in roses and will facilitate future breeding of this important ornamental crop.

## Results

### Haplotype-resolved genome assembly for the tetraploid modern rose

To develop a reference genome for tetraploid modern roses, we sequenced the genome of cultivar ‘Samantha’ (2n = 4x = 28), which had an estimated tetraploid genome size of approximately 2.46-2.55 Gb based on the flow cytometry analysis (Supplementary Fig. 1 and Supplementary Table 1). We generated 84 Gb of PacBio HiFi reads covering approximately 34× of the ‘Samantha’ genome, 89 Gb of Illumina paired-end reads, and 521 Gb of chromosome conformation capture (Hi-C) sequences. The final assembled genome contained 36,850 contigs, with a total size of 2,698 Mb and an N50 length of 188 kb. The Hi-C data was used to further assemble 93.7% of the contigs (2,519 Mb) into 28 chromosomes (Supplementary Fig. 2 and Supplementary Table 2).

Genome resequencing data of 62 accessions of original species were used to assign chromosomes into four haplomes (four set of chromosomes, each with seven chromosomes). We named the haplome closest to the Chinese original species as ChrA, and the haplome closest to the European original species as ChrB. ChrC and ChrD also originate from Chinese and European original species, respectively (Supplementary Fig. 3). ChrA, ChrB, ChrC, and ChrD had total lengths of 641 Mb, 593 Mb, 676 Mb, and 607 Mb, respectively (Supplementary Table 3). Around 95% of the RNA-Seq reads from leaves, floral buds, and petals could be aligned to each haplome of the assembly. Meanwhile, BUSCO^8^ evaluation showed that 94.0-96.4% of the core genes were completely captured in each of the four haplomes, and the entire haplotype-resolved genome captured 99.3% of the BUSCO genes, with the majority in four copies (Supplementary Table 4 and Supplementary Fig. 4). Furthermore, the LTR Assembly Index^9^ (LAI) for each of the four haplomes ranged from 13.48 to 14.9. Similarly, k-mer analysis showed the high completeness and highly phased nature of the ‘Samantha’ genome assembly (Supplementary Fig. 5). Finally, the integrity of the chromosome-scale assembly of ‘Samantha’ was confirmed using the phased genetic map of tetraploid modern rose^10^ (Supplementary Fig. 6). Together, these results supported the high accuracy and completeness of the ‘Samantha’ genome assembly.

The four haplomes showed different divergence times from the *R. chinensis* ‘Old Blush’, which were inferred from the distributions of pairwise synonymous substitution rates (*K*s) of orthologous genes (Supplementary Fig. 7), with ChrA having the smallest *Ks* value and ChrB the largest. This further confirmed the accuracy of haplome and subgenome classification. The four haplomes of ‘Samantha’ showed high collinearity, as well as to the genome of ‘Old Blush’ (Fig. 1 and Supplementary Fig. 8).

**Fig. 1.**
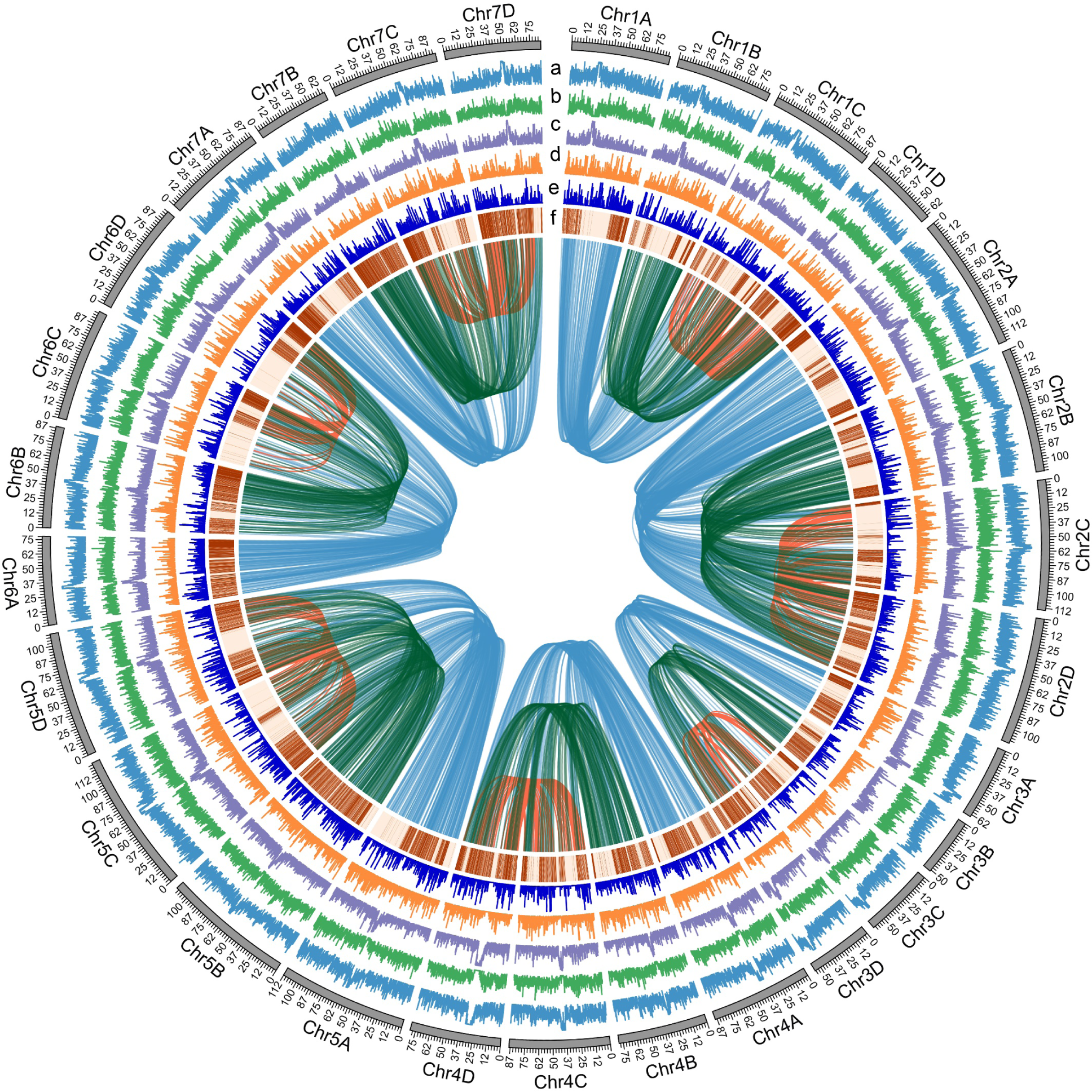
Genomic landscape of *Rosa hybrida* ‘Samantha’. **a-f**, LTR density (**a**), *Copia*-type LTR-RT density (**b**), *Gypsy*-type LTR-RT density (**c**), DNA transposon density (**d**), gene density (**e**), and gene expression levels (**f**) in 500 kb non-overlapping windows. Syntenic blocks between ChrA, ChrB, ChrC and ChrD are shown at the center.

A total of 59.86% of the ‘Samantha’ genome assembly was identified as repeat sequences, with the majority (45.85% of the assembly) annotated as long terminal repeat retrotransposons (LTR-RTs) (Supplementary Table 5). A total of 139,617 protein-coding genes were predicted from the genome, of which 132,795 (95.11%) were functionally annotated (Supplementary Table 6). The four haplomes harbored a total of 131,295 genes, with 33,224, 32,144, 33,218, and 32,709 in ChrA, ChrB, ChrC and ChrD, respectively.

### Subgenome features of the modern rose

In an allopolyploid genome, it is common for one parental subgenome to retain more genes, exhibit higher gene expression, and/or experience stronger purifying selection than other subgenome(s). This phenomenon is known as subgenome dominance^11^. The four haplomes of ‘Samantha’ were found to have similar gene contents. The GC content, gene structure, and repetitive element distribution did not differ significantly between the four haplomes (Supplementary Table 7). Gene family clustering analysis revealed that the number of unique gene families among the four haplomes was similar. KEGG annotation revealed that ChrA had unique gene families enriched with those involved in steroid biosynthesis, ChrB in phenylalanine, tyrosine, and tryptophan biosynthesis, ChrC in amino acid biosynthesis, and ChrD in sulfur metabolism (Supplementary Fig. 9 and Supplementary Table 8).

To understand the evolution of the tetraploid modern rose genome after genome merger by hybridization, we investigated the homoeologous gene contents retained in the two subgenomes, using ChrA and ChrB as the showcase. No chromosome-level gene loss bias was found between ChrA and ChrB, although bias in some local regions was observed (Fig. 2a). Consistent with no whole-genome bias in gene loss, all seven chromosomes in ChrA had similar *Ka* (nonsynonymous substitution rate) to *Ks* ratios compared to their homoeologous chromosomes in ChrB (Fig. 2b and Supplementary Table 9), supporting that the two subgenomes have evolved symmetrically and have been under a similar strength of purifying selection.

**Fig. 2.**
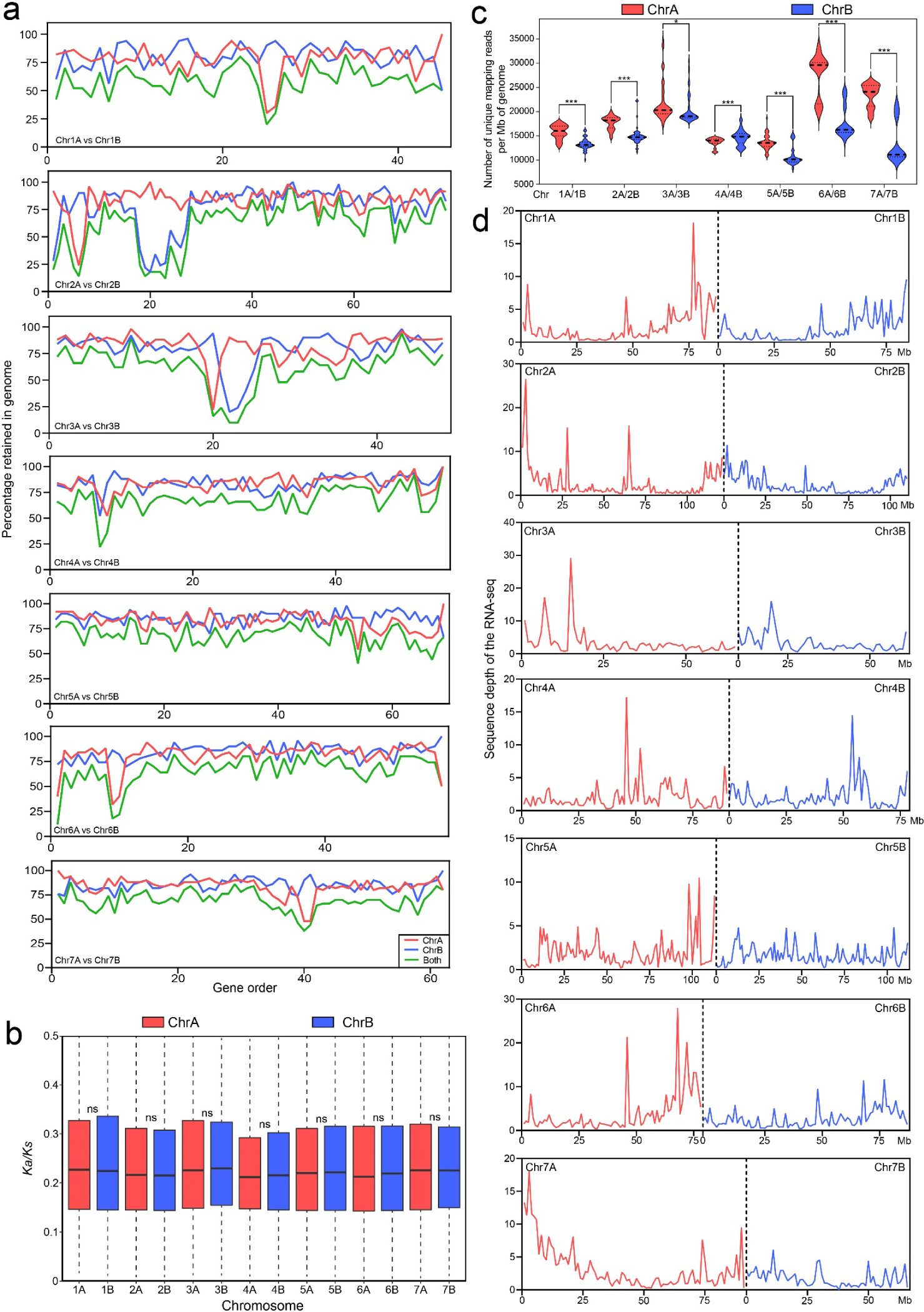
Subgenome features of *R. hybrida* ‘Samantha’. **a,** Gene retention between the two subgenomes of ‘Samantha’. The percentages of genes retained in ChrA (red), ChrB (blue), and both subgenomes (green) along chromosomes are shown. **b,** Boxplot of the *Ka*/*Ks* distribution of homoeologous genes in chromosomes of ChrA (red) and ChrB (blue). The subgenomes of ‘Samantha’ were used to construct orthologous gene pairs with the strawberry genome. **c,** Average number of uniquely mapped reads in 54 RNA-Seq samples across the two subgenomes. **d,** Distribution of the average RNA-Seq read coverage depth of 54 samples across the two subgenomes.

RNA-Seq data from 54 ‘Samantha’ samples were used to detect expression dominance between the subgenomes. Uniquely mapped reads revealed that expression biases existed between subgenomes. Overall, the Chinese-origin chromosomes had a higher expression, with the exception of Chr4 (Fig. 2c and 2d). We detected 6,756 syntenic tetrads (27,024 genes) (one gene in each of the four haplomes) in the ‘Samantha’ genome (Supplementary Table 10). Among them, 4,574 (67.7%) were found to maintain a balanced expression trend, with 3.4% of tetrads possessing a single dominant expressed gene (Supplementary Fig. 10). Therefore, the bias in gene expression between subgenomes is not likely due to the differences in expression of syntenic tetrads.

### Genomic variation map

We performed genome resequencing of 214 accessions in the *Rosa* genus, including wild accessions from different sections of the genus (Supplementary Notes 1-2), intermediate old cultivars (bred from hybridization among old cultivars from China and Europe), and modern cultivars (Supplementary Table 11). Together with ‘Samantha’ and another 18 accessions with publicly available resequencing data^6^, a total of 233 accessions were analyzed in this study, including 62 (wild species and old cultivars) from section Chinenses, 16 from section Rosa, 18 from section Synstylae, 11 from section Cinnamomeae, 4 from section Caninae, 15 from other sections (including Pimpinellifoliae, Microphyllae, Banksianae, Bracteatae, and Laevigatae), 65 intermediate cultivars, and 42 modern cultivars. In total, we identified 14,060,480 SNPs, among which 10,471,525 (74.47 %) were identified within Chinenses, 8,975,644 (63.84%) within Rosa, and 9,985,128 (71.02%) within Synstylae. As for genomic locations, 9,514,445 (67.67%) were in intergenic regions, 2,889,889 (20.55%) in intron regions and 1,617,063 (11.50%) in coding regions. Among SNPs in coding regions, 837,643 were nonsynonymous and 760,596 were synonymous (Supplementary Table 12). Furthermore, a total of 17,301 SNPs introduced premature stop codons in 11,327 genes. Among these nonsense SNPs, we identified one in the *Centroradialis* (*CEN; RhySMT6AG254700*) gene that encodes a protein homologous to Arabidopsis TERMINAL FLOWER 1 (TFL1), a key regulator of flowering time and inflorescence development^12, 13^. This SNP mutation (C->T) was highly conserved in wild species including those from sections Synstylae, Cinnamomeae, Pimpinellifoliae, Microphyllae, Bracteatae, and Banksianae, while in modern cultivars, the majority had the homozygous C/C or heterozygous C/T genotype at this locus (Supplementary Fig. 11a), which could lead to the predominance of solitary flowers in modern cultivars. Furthermore, we found that among the 233 rose accessions, this variant was significantly associated with the inflorescence type (Supplementary Fig. 11b and Supplementary Table 13), providing a valuable marker for future modern rose breeding.

We then evaluated the nucleotide diversity of different rose populations. The nucleotide diversity (π) of 10 different wild sections ranged from 1.12×10^-3^ to 2.94×10^-3^. The intermediate cultivars showed the highest degree of nucleotide diversity (3.79×10^-3^), also with 99.44% of the SNPs being detected in this group. The nucleotide diversity of modern cultivars displayed a slightly decreased level (3.49×10^-3^) compared to that of the intermediate cultivars, but significantly higher than that of any of the wild groups. This suggests a weak bottleneck effect during the domestication of modern roses and indicates that modern roses still have great genetic potential for improvement.

### Origin of modern rose

According to historic documents and genetic analysis, 8-20 species of *Rosa* are thought to be involved in modern rose creation^4^, though the exact donors are still in dispute. To better understand the origin and breeding history of modern rose as a new population, a phylogenetic tree of the 233 accessions was constructed using high-quality SNPs (Fig. 3a). Modern cultivars were located within the clade comprising all members of Chinenses and part of Synstylae, suggesting possible major contributions of Chinenses and Synstylae species to modern cultivars. Intermediate cultivars, as early hybrids, were mainly distributed around modern cultivars on the phylogenetic tree. Consistently in the principal component analysis (PCA), accessions from section Cinnamomeae and group Other were clearly separated from modern cultivars (Fig. 3b). The intermediate group exhibited the broadest diversity. By contrast, modern cultivars formed a tighter cluster within the intermediate group, indicating a narrower genetic variation after human selection along continuous breeding history. The closer distribution of modern cultivars to Chinese accessions (section Chinenses and part of Synstylae) was consistent with the increased Chinese/European allele ratio discovered in cultivars bred in 18^th^-19^th^ century^14^.

**Fig. 3.**
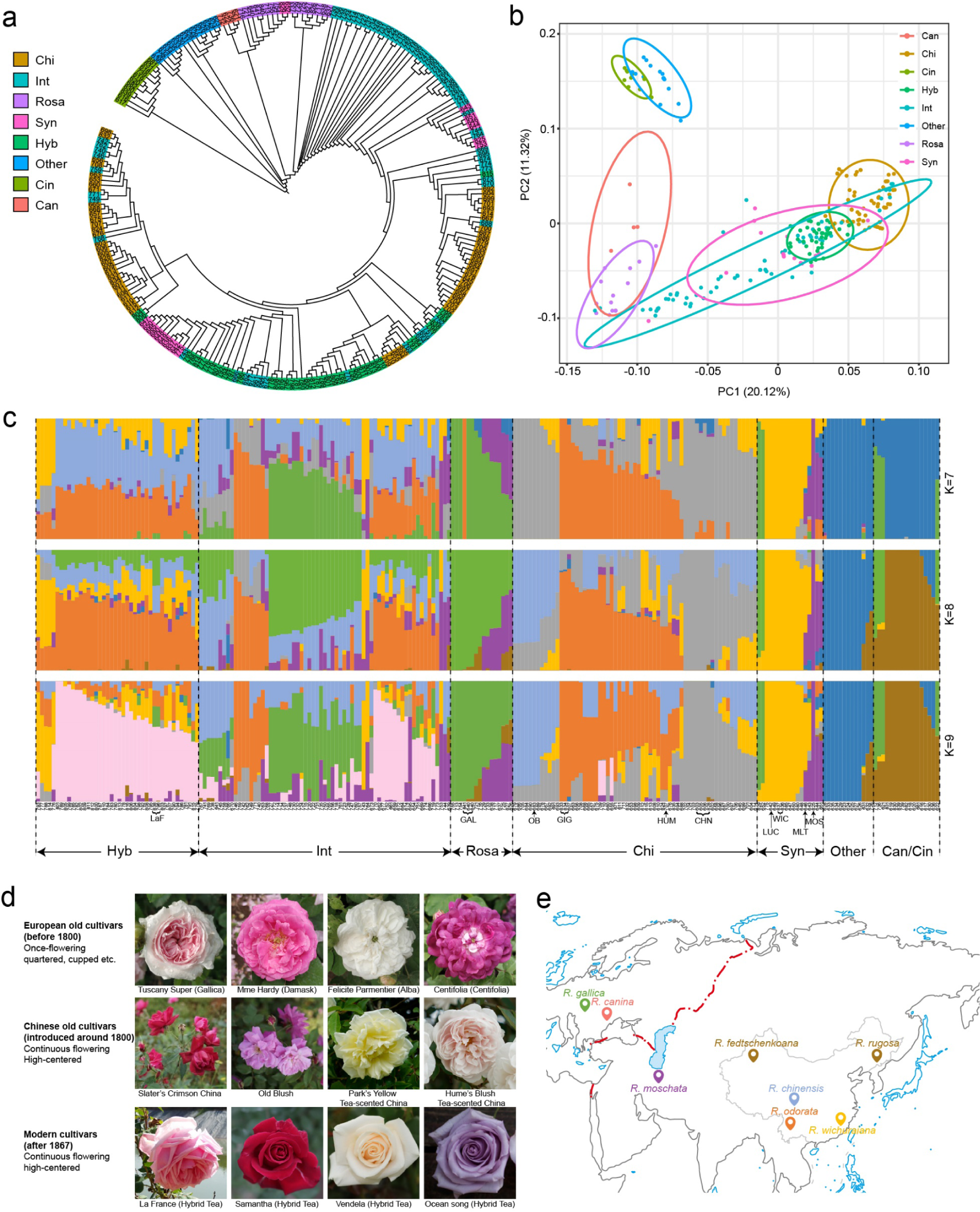
Population structure of 233 accessions within genus *Rosa*. **a,** Phylogenetic tree constructed using high-quality SNPs. Hyb, modern cultivars, *R. hybrida*; Int, intermediate cultivars; Syn, section Synstylae; Chi, section Chinenses; Rosa, section Rosa; Can, section Caninae; Cin, section Cinnamomeae; Other, sections Pimpinellifoliae, Microphyllae, Bracteatae, and Banksianae. **b,** Principal component analysis of the 233 accessions. **c,** Bayesian model-based clustering of the 233 accessions with the number of ancestry kinships (*K*) from 7 to 9. Each vertical bar represents an accession. Each color represents one putative ancestral background and the y axis quantifies ancestry contribution. LaF, first Hybrid Tea rose cultivar ‘La France’; HUM, *R. odorata* ‘Hume’s Blush Tea-scented China’; OB, *R. chinensis* ‘Old Blush’; WIC, *R. wichuraiana*; MLT, *R. multiflora*; LUC, *R. luciae*; MOS, *R. moschata*; GIG, *R. odorata* var. gigantea; CHN, *R. chinensis* var. spontanea; GAL, *R. gallica.* **d,** Changes of flower shape along the rose breeding history. **e,** Geographical distribution of the eight original species. *R. chinensis* is native to Zhejiang, Guangdong, Guangxi, Fujian, and Taiwan in China. *R. wichuraiana* is native to Hubei, Sichuan, and Guizhou in China. *R. rugosa* is native to northern China, Japan, and Korea. *R. gallica* is native to central and southern Europe and Western Asia. *R. canina* is native to Europe and Western Asia. *R. fedtschenkoana* is native to Xinjiang in China. *R. odorata* is native to Yunnan in China. *R. moschata* is believed to have originated from South Asia, including Iran, Afghanistan, Pakistan, and India.

To infer possible progenitors of modern roses, we further performed population structure analysis (Fig. 3c). Our Δ*K* analysis revealed that seven populations (*K*=7) were optimal for these 233 accessions, while nine populations (*K*=9) were also close to optimal (Supplementary Fig. 12). At *K*=7, accessions from section Rosa were quite distinguishable from other accessions and they genetically contributed to part of intermediate cultivars rather than modern cultivars. At *K*=8, *R. chinensis* ‘Old Blush’ was separated from the original wild species *R. chinensis* var. *spontanea* and was quite genetically different from ‘Hume’s Blush Tea-scented China’, which was derived from wild species *R. odorata* var. *gigantea*. For modern cultivars, they mainly harbored genetic compositions from sections Chinenses, Rosa, and Synstylae. We found that, ‘Hume’s Blush Tea-scented China’, a representative of cultivars derived from *R. odorata* var. *gigantea*, contributed more to the genetic composition of modern cultivars than ‘Old Blush’. At *K*=9, modern roses showed similar genetic components among each other even with complex origins. This may be the result of extensive hybridization and human selection made by numerous breeders and masses of rosarians and amateurs worldwide. Considering that favored mutations can be selected and well preserved through clonal propagation in rose, it is not surprising that modern rose cultivars have developed from several heterogeneous ancestors into a genetically relatively similar group.

Within modern roses, cultivars with larger genetic contributions from section Synstylae (Fig. 3c) had flowers in large clusters or trusses and thus classified as group Floribunda (Supplementary Note 3). It has long been recognized that genome introgression from *R. multiflora* into modern cultivars has passed down its typical inflorescence trait (flowers in clusters) to Floribunda, supported by morphological evidence (Supplementary Fig. 13). Genetic structure of all the three wild species from section Synstylae, *R. multiflora*, *R. luciae* and *R. wichuraiana*, could not be further distinguished from each other (Fig. 3c), and thus the exact contribution of each species to this inflorescence trait of modern roses still remained unclear.

Before the introduction of Chinese germplasms around 1800, European old cultivars, e.g., Gallica, Damask, Alba, and Centifolia (top panel in Fig. 3d), were generally once-flowering and had a relatively similar quartered or cupped flower shape. Relatively detailed records of introduced Chinese accessions focused on the four well-known cultivars (middle panel in Fig. 3d). Continuous flowering is a highly valued ornamental trait and *R. chinensis* ‘Old Blush’ exhibiting this feature is always believed to be an important Chinese progenitor to modern roses^1, 6, 7^. Unexpectedly, its contribution to modern roses at the genomic level seemed not large (Fig. 3c). In addition to the continuous-flowering trait of the four introduced Chinese old cultivars, the dark red color of *R. chinensis* ‘Slater’s Crimson China’ and the distinct tea scent from two other *R. odorata* cultivars (‘Hume’s Blush Tea-scented China’ and ‘Park’s Yellow Tea-scented China’) made them all extensively used in hybridizations for modern rose formation. The two *R. odorata-*derived cultivars are believed to only contribute to the tea scent trait, which has been lost in most modern rose cultivars. Interestingly, our population structure analysis suggested that they contributed the most to the genetic composition of modern cultivars (*K*=7-8 in Fig. 3c) and may thus laid the foundation of the favored flower shape of modern cultivars (bottom panel in Fig. 3d). It is worth mentioning that *R. odorata*-derived old cultivars exhibit a high-centered flower shape distinct from that of European old cultivars, and their petal edges can be reflexed backward at later opening stages. The flower shape of modern rose cultivars (bottom panel in Fig. 3d) mimicked that of *R. odorata*-derived old cultivars, while with increased petal numbers. Collectively, *R. odorata* and derived cultivars contributed a larger portion to the genome structure and favored flower shape of modern cultivars than *R. chinensis* ‘Old Blush’ and *R. gallica* (Fig. 3c, d). Ultimately, we clarified the contribution of six original species to modern cultivars, and the geographical distribution^15^ of these original species highlights the importance of artificial hybrid breeding (Fig. 3e).

### Genomic composition of *R. hybrida* ‘Samantha’

The complex hybrid history of modern roses makes it difficult to identify the origins of their chromosomes. Population structure analysis reveals six potential original ancestors of modern roses (Fig. 3c). In order to better analyze the genomic composition of *R. hybrida* ‘Samantha’, the origins of different regions of each chromosome were inferred based on the mapped read depth and the homozygous variant rate of the wild original species. The consensus results of the two approaches were used to infer possible ancestral origins of regions in each chromosome (Fig. 4a, b and Supplementary Fig. 14). We inferred that the largest proportion of ‘Samantha’ genomes is derived from the *R. odorata*, accounting for 17.42%, and *R. chinensis*, *R. wichuraiana*, *R. gallica*, *R. moschata* and *R. rugosa* contribute 10.08%, 9.74%, 9.42%, 7.58% and 5.8% of the genome, respectively (Fig. 4c). Interestingly, the ‘Samantha’ chromosomes exhibited a high degree of chimerism, each derived from multiple ancestral species except Chr7B, indicating extensive homeolog exchanges and/or introgressions in the ‘Samantha’ genome (Supplementary Fig. 14) and consistent with the segmental allotetraploid nature of modern rose^10^. Chr7B was entirely derived from *R. rugosa* (section Cinnamomeae) and was notably shorter than the other three homologous chromosomes, which we speculate might be the reason for the absence of homeolog exchanges. Furthermore, a large inversion between the two subgenomes was observed on chromosome 7 (Fig 4d). Genomic regions of Chr7A and Chr7C (subgenome A) harboring this inversion were both derived from *R. odorata*, and those of Chr7B and Chr7D (subenome B) were derived from *R. rugosa*, and *R. wichuraiana*, respectively. Further collinearity analysis with the three published *Rosa* genomes, including *R. chinensis* ‘Old Blush’, *R. wichuraiana*, and *R. rugosa*, revealed the same inversion between subgenome A and all these *Rosa* genomes (Supplementary Fig. 15). Notably, the KEGG and GO analysis showed that genes in the inverted regions were enriched with those involved in glycerophospholipid metabolism and membrane organization, which play important roles in maintaining cell membrane stability and permeability, and enhance plant stress tolerance, growth, and development.

**Fig. 4.**
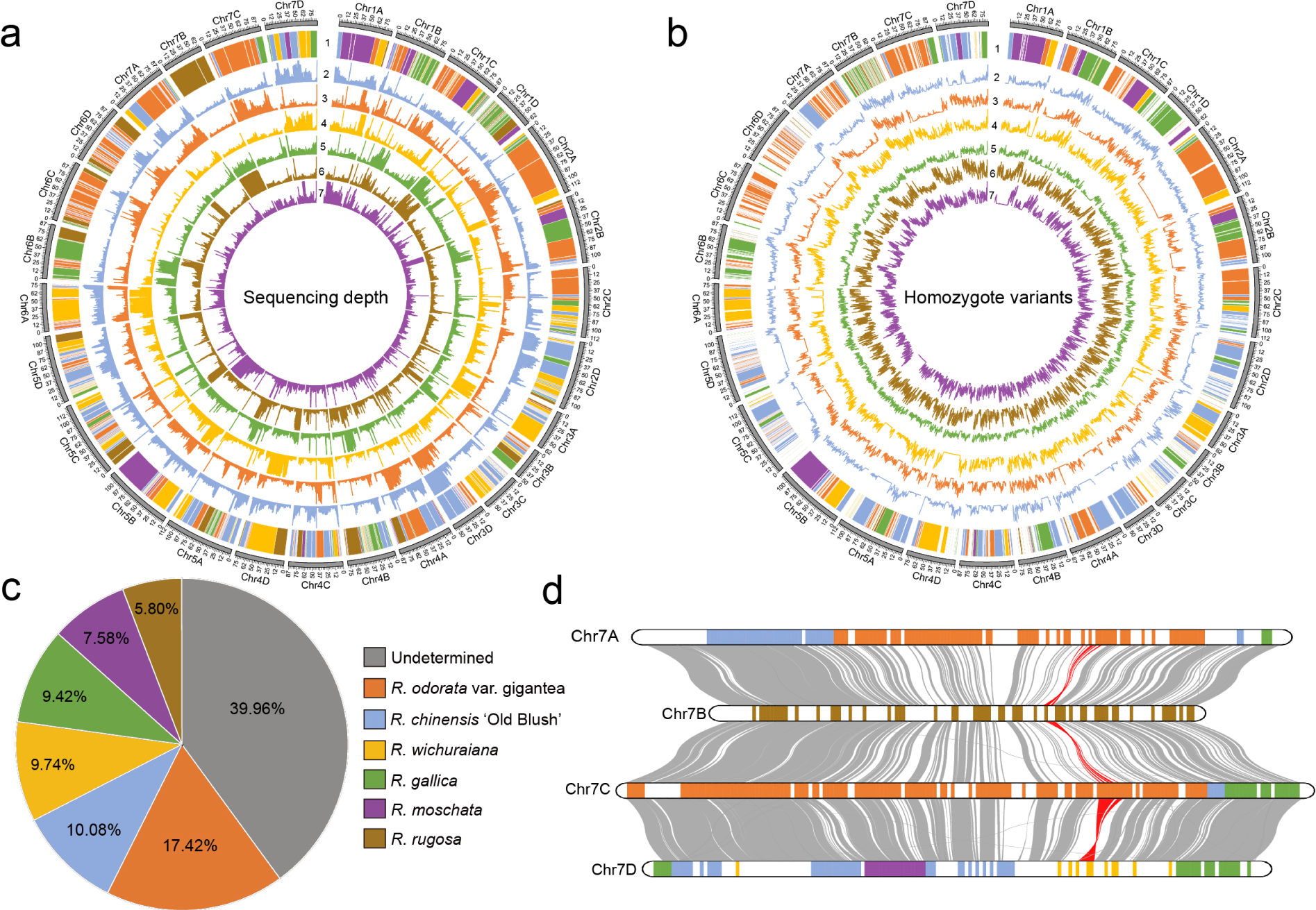
Genetic compositions of the ‘Samantha’ chromosomes. **a,** Origins of different regions on the chromosomes (1) inferred by the depth coverage of reads from wild ancestors, *R. chinensis* ‘Old Blush’ (2), *R. odorata* var. *gigantea* (3), *R. gallica* (5), *R. wichuraiana* (4), *R. rugosa* (6), and *R. moschata* (7). **b,** Origins of different regions on the chromosomes (1) inferred by the homozygote variant rate between ‘Samantha’ and wild ancestors, *R. chinensis* ‘Old Blush’ (2), *R. odorata* var. *gigantea* (3), *R. gallica* (5), *R. wichuraiana* (4), *R. rugosa* (6), and *R. moschata* (7). **c,** Summary statistics of contributions of the six wild ancestors to the ‘Samantha’ genome. **d,** Collinearity of the four haplomes of chromosome 7. Colors on the chromosomes represent origins from different original species, *R. chinensis* ‘Old Blush’ (blue), *R. odorata* var. *gigantea* (orange), *R. gallica* (green), *R. wichuraiana* (yellow), *R. rugosa* (brown), and *R. moschata* (purple). The inversion is shown in red.

### Artificial selection during modern rose breeding

To investigate genomic regions that have been under selection during rose breeding, we conducted selective sweep analysis between modern cultivars and intermediate old cultivars. Considering that the ploidy level could interfere with this analysis, we chose cultivars confirmed to be tetraploid from intermediate cultivars (referred as Int_4) and modern cultivars (referred as Hyb_4) for this analysis (Supplementary Table 11). Int_4 had a higher nucleotide diversity (π = 3.87×10^-3^) than that of Hyb_4 (π = 3.51×10^-3^), reflecting the function of human selection during cultivar improvement (Fig. 5a). Additionally, Int_4 exhibited a faster LD decay than Hyb_4 (Fig. 5b), again supporting the decreased genetic diversity for recently formed modern cultivars. Through comparing Int_4 with Hyb_4 and using combined filtering of π ratio and *F*_ST_, selective sweep regions were identified with a total length of 56.69 Mb, which contained 3,411 genes (Supplementary Table 14). Within these regions, genes related to continuous flowering, floral organ development, flower color, senescence, growth, and disease resistance were revealed (Fig. 5c).

**Fig. 5.**
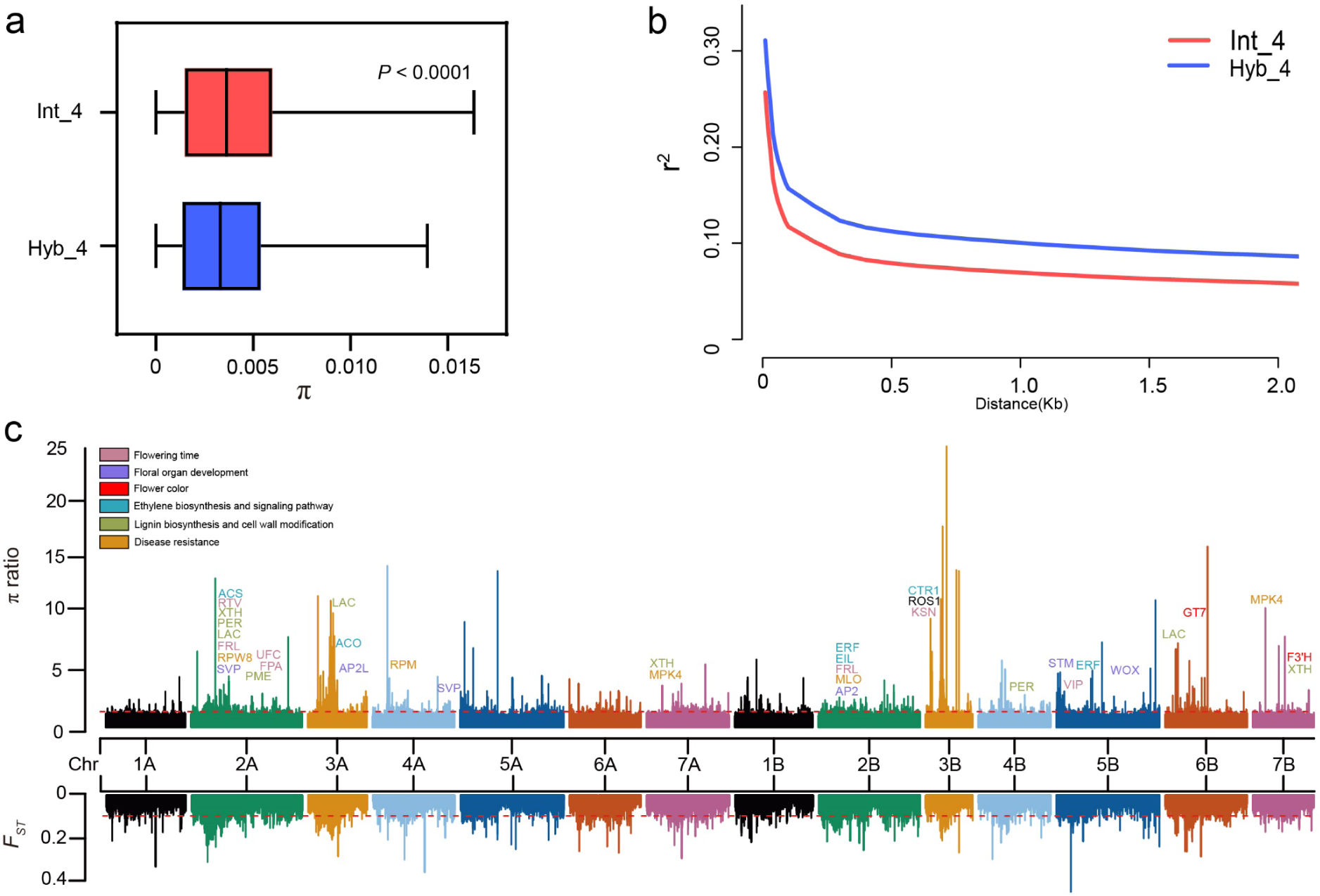
Genome-wide distribution of selective sweeps in *R. hybrida*. **a,** Nucleotide diversity (π) of tetraploid intermediate (Int_4; n = 45) and tetraploid modern rose cultivars (Hyb_4; n = 32). For each box plot, the lower and upper bounds of the box indicate the first and third quartiles, respectively, and the center line indicates the median. **b,** LD decay in Int_4 and Hyb_4. **c,** π ratio and *F*_ST_ analyses between Int_4 and Hyb_4 reveal potential selective regions across rose chromosomes. Horizontal red dashed lines indicated the top 5% cutoff values. Gene abbreviations: *KSN*, *Koushin*; *UFC*, *upstream of FLC*; *FPA*, *flowering time control gene FPA*; *FLK*, *FLOWERING LOCUS KH DOMAIN*; *FRL*, *FRIGIDA-like*; *VIP*, *VERNALIZATION INDEPENDENCE*; *GT7*, *UDP-glucose flavonoid 3-O-glucosyltransferase 7*; *RTV*, *RELATED TO VERNALIZATION1*; *SVP*, *SHORT VEGETATIVE PHASE*; *AP2*, *APETALA2*; *AP2L*, *APETALA2-like*; *F3’H*, *flavanone 3-hydroxylase*; *CTR1*, *CONSTITUTIVE TRIPLE RESPONSE 1*; *ACS*, *1-aminocyclopropane-1-carboxylate synthase*; *ACO*, *1-aminocyclopropane-1-carboxylate oxidase*; *EIL*, *ethylene-insensitive like*; *ERF*, *ethylene response factor*; *LAC*, *LACCASE*; *PER*, *PEROXIDASE*; PME, *pectinesterase*; *XTH*, *xyloglucan endotransglucosylase/hydrolase*; *STM*, *SHOOT MERISTEMLESS*; *WOX*, *WUSCHEL related homeobox*; *RPW8*, *resistance to powdery mildew 8*; *MLO*, *mildew resistance locus O*; *RPM*, *disease resistance gene RPM*; *MAPK4*, *mitogen-activated protein kinase 4*.

The ability of continuous flowering is among the most important traits of modern roses. It is the basis of cut rose production and also highly valued for garden/potted roses. This character is believed to be inherited from *R. chinensis* cultivars^1, 6^. The *Koushin* (*KSN*) gene in rose has been found to be involved in the determination of once/continuous flowering^16, 17^. Here, the *KSN* gene (*RhySMT3BG120200*) was found in a selective sweep region (Supplementary Fig. 16a-b). Clustering analysis of SNP genotype in this selective sweep region further supported the contribution of section Chinenses to the continuous flowering of modern roses (Supplementary Fig. 16c). Furthermore, genes encoding regulators involved in vernalization^18^ (e.g., *VERNALIZATION INDEPENDENCE*, *FRIGIDA-like*, *RELATED TO VERNALIZATION1*) and the autonomous pathway^19^ (e.g., *SHORT VEGETATIVE PHASE*, *flowering time control protein FPA*) were also intensively selected, suggesting that the trait of continuous flowering may have been realized by fixing genes in separate flowering pathways.

Flower meristem identity genes *SHOOT MERISTEMLESS* (*STM*) and *WUSCHEL related homeobox* (*WOX*) were found to have undergone selection during modern rose breeding. They participate in the formation of the shoot apical meristem and affect the development of floral organs. The number of petals is another important ornamental trait and has been increased greatly along modern rose breeding. In rose, floral identity genes *APETALA 2* (*AP2*) and *APETALA 2-like* (*AP2L*) have been proven to regulate petal numbers through stamen petaloidy^20–22^. Here, we identified that *AP2* (*RhySMT2BG131100*) and *AP2L* (*RhySMT3AG121100*) have been under selection during modern rose breeding, resulting in reduced nucleotide diversity in modern cultivars (Supplementary Fig. 17a-b, d-e). We performed clustering on the SNP genotypes of these two selective sweep regions that contained *AP2* and *AP2L*, respectively, and found that SNP profiles of modern cultivars were highly similar to those of section Chinenses (Supplementary Fig. 17c, f). This directly demonstrates the crucial contribution of section Chinenses to the petal number in modern roses. Moreover, petal color is among the most attractive characteristics of modern roses. *UDP-glucose flavonoid 3-O-glucosyltransferase 7* (*GT7*; *RhySMT6BG069000*) and *flavanone 3-hydroxylase* (*F3’H*; *RhySMT7BG191700*) involved in floral pigment biosynthesis^23^ were also identified in selective sweep regions.

Opening and senescence of flowers are essential parameters determining the quality of modern roses. Ethylene has been extensively documented to play a pivotal role in governing flower opening and senescence in rose^24–26^. Ethylene-related genes were found in several highly selected regions, including biosynthesis-related genes (*ACO*, *ACS*) and genes encoding signaling components (*CTR1, EIL*) and ethylene-responsive transcription factors (*ERFs*), indicating that both ethylene biosynthesis and signaling pathways have been selected during the breeding history of modern roses. *CTR1*^27^, the key component of the ethylene signal transduction pathway, has been under selection (Supplementary Fig. 18a-b), and its SNP genotype profile was similar to that of *R. chinensis* ‘Semperflorens’ (Supplementary Fig. 18c). Furthermore, *ROS1*^28^, encoding a DNA demethylase, was also found in this selective sweep region (Supplementary Fig. 18a), indicating that DNA methylation might play a critical role in determining rose traits during its breeding.

Additionally, genes encoding enzymes related to lignin biosynthesis and cell wall modification, e.g., *xyloglucan endotransglucosylase/hydrolase*^29^ (*XTH*), *LACCASE* (*LAC*), *PEROXIDASE* (*PER*), and *pectinesterase* (*PME*), were found in selective sweep regions. Mutations in *LAC* and *PER* could result in defects in lignification and thus growth arrest^30, 31^. *PMEs* are found to be associated with plant height in soybean^32^ and fruit weight in peach^33^. Therefore, selection of these genes in modern roses might be related to growth rate and yield. In modern rose breeding, selection for disease resistance has always been a main target^34^. Therefore, it is not surprising that compared to intermediate cultivars, disease resistance in modern cultivars has been selected during the last two hundred years of modern rose breeding history. Two genes homologous to Arabidopsis *MAPK4* that plays a crucial role in regulating plant innate immune responses^35^, one gene homologous to RPM conferring resistance to bacterial pathogens^36^, one gene homologous to Arabidopsis *RPW8* that is involved in powdery mildew resistance^37^, and a rose *mildew resistance locus o* (*MLO)* gene, also responsible for powdery mildew resistance^34, 38^, were identified in the selective sweeps. The selection of these genes further confirmed the importance of disease resistance in breeding of modern cultivars.

## Discussion

The modern rose, which is mostly tetraploid, is a hybrid resulting from the crossbreeding of multiple ancestors from both eastern and western of Eurasia. It has inherited important agronomic traits from its European and Chinese progenitors. While one of the Chinese diploid progenitor *R. chinensis* ‘Old Blush’ has been sequenced recently^6, 7^, a reference genome of the tetraploid modern rose is of irreplaceable significance for understanding its origin and domestication, as well as for improving this important ornamental crop. In this study, we present a haplotype-resolved genome assembly of the tetraploid modern rose cultivar ‘Samantha’, along with a comprehensive genome variation map of 233 accessions from wild species and modern and old cultivars in the *Rosa* genus. This work allows us to reconstruct the history of divergence, hybridization, and domestication of roses. The genome of ‘Samantha’ is one of the few polyploid plant genomes reported to date, providing a powerful resource for future comparative and evolutionary genomic studies.

Population genomic analyses reveal that old Chinese and European cultivars originated from reticulate evolution among several wild species within *Rosa*. *R. odorata*, in addition to *R. chinensis* ‘Old Blush’^3, 14^, played an important role as the Chinese progenitor of modern rose during the creation of modern rose through hybridizations in the 19^th^ century. Notably, some previously unappreciated wild species, such as those in section Synstylae, have also played important roles in the formation of modern rose, particularly the Floribunda type with clustered flowers.

Over thousands of years, the cultivation and domestication of old cultivars in both China and Europe have accumulated a significant number of ideal agronomic traits. These traits have been retained and further selected in modern rose, making it one of the most important ornamental crops worldwide. To identify genome regions under selection during modern rose breeding that are potentially associated with important agronomic traits, such as continuous flowering, fragrance, senescence and disease resistance, we compared modern roses with old cultivars. Continuous flowering is a core trait of modern rose cultivars, and therefore selection of related genes is expected. On the other hand, extending the ornamental period of modern roses is important, especially for cultivars used for cut flowers. It is important to develop cultivars that have low sensitivity to ethylene in order to delay senescence. By controlling the crucial component upstream of the ethylene signaling cascade, it is possible to effectively reduce ethylene signal transduction, thereby minimizing the ethylene response and extending the life of cut flowers. The absence of a rich scent in a large number of modern cultivars is an unexpected trait overlooked during breeding. Our study did not identify any genes related to scent-synthesis in genome regions under selection. Since the domestication and breeding of roses have only occurred for a few hundred years, we believe that scent could become a new targeted trait in future rose breeding to meet consumers’ preference.

Despite the short history of modern rose, frequent crossing and backcrossing make it a unique group. In this study, we present new insights into the structure and evolution of the tetraploid modern rose genome, nucleotide diversity, and fixation of horticultural traits. Together with the haplotype-resolved high-quality modern rose genome and single-base resolution variation map, the findings of this study provide valuable information for future research on the agronomically important traits of roses, and facilitate marker-assisted and genomic selection-based breeding in the future.

## Methods

### Sample preparation and sequencing

The widely cultured modern rose cultivar, ‘Samantha’, which produces medium-sized red flowers and dark green, leathery foliage, was selected for reference genome sequencing. PacBio HiFi libraries with inserts of ∼15 kb were prepared following the PacBio HiFi library construction protocol, and sequenced on the PacBio Sequel II system. For Illumina sequencing, a DNA sequencing library was constructed with an insert size of ∼250 bp and sequenced on an Illumina NovaSeq 6000 platform. Hi-C library was prepared following the method described in Belton et al.^39^ with an improved modification. Briefly, calluses were vacuum infiltrated in nuclei isolation buffer supplemented with 2% formaldehyde for 15 min and then ground with liquid nitrogen and washed orderly by extraction buffer Ⅰ (0.4 M sucrose, 10 mM Tris-HCl, pH 8.0, 10 mM MgCl_2_, 5 mM β-mercaptoethanol, 0.1 mM phenylmethylsulfonyl fluoride [PMSF] and protease inhibitor), extraction buffer Ⅱ (0.25 M sucrose, 10 mM Tris-HCl, pH 8.0, 10 mM MgCl_2_, 1% Triton X-100, 5 mM β-mercaptoethanol, 0.1 mM PMSF and protease inhibitor) and extraction buffer Ⅲ (1.7 M sucrose, 10 mM Tris-HCl, pH8.0, 0.15% Triton X-100, 2 mM MgCl_2_, 5 mM β-mercaptoethanol, 0.1 mM PMSF and protease inhibitor). Overnight digestion was applied with restriction enzyme (400 units MboI) at 37 ℃ on rocking platform. The following steps were conducted as in Belton et al.^39^. The Hi-C libraries were sequenced on an Illumina HiSeq platform.

A total of 214 *Rosa* accessions representing wild species, intermediate cultivars and modern cultivars were selected for genome resequencing. High-quality genomic DNA was extracted from young fresh leaves using the cetyltrimethylammonium bromide (CTAB) method. Illumina paired-end libraries with insert sizes of ∼250 bp was constructed using the Illumina Genomic DNA Library Prep Kit following manufacturer’s recommendations and sequenced on an Illumina NovaSeq 6000 platform.

### Genome size estimation and ploidy analysis

The genome size of ‘Samantha’ was estimated using flow cytometry^40^. Leaves were placed in a 500 μL CyStain PI Absolute P Nuclei Extraction Buffer, chopped with a razor blade, and then filtered through a 50-μm filter. The collected cells were combined with 2,000 μL of CyStain PI Absolute P Staining Buffer and incubated in the dark for 30 min. The nuclei suspension was analyzed using CyFlow Space Flow Cytometer (Sysmex Partec). Genome size estimations were independently determined using *Solanum pimpinellifolium* and *R. chinensis* ‘Old Blush’ as the internal reference standards.

Ploidy analysis was conducted with young fresh leaf tissue, using the same cell extraction approach described above for the genome size estimation. CyStain UV Precise P Staining Buffer was added, and samples were incubated at room temperature for 1 min. The ploidy level of each sample was analyzed using CyFlow Ploidy Analyser (Supplementary Fig. 19).

### RNA extraction and transcriptome sequencing

Total RNA of floral buds and petals from three different developmental stages, 0, 1, and 2 according to Ma et al.^41^, was extracted using the hot borate method as previously described^26^. Strand-specific RNA-Seq libraries were constructed using the protocol described in Zhong et al.^42^ and sequenced on the Illumina HiSeq 2000 system. Three biological replicates were conducted for each sample.

### *De novo* genome assembly of ‘Samantha’

PacBio HiFi reads of ‘Samantha’ were *de novo* assembled into contigs using HiCanu^43^ (v.2.0) with default parameters. Potential contaminations from microorganisms were then detected in the assembly by aligning the assembled contigs to NCBI non-redundant nucleotide database using BLASTN^44^ (v.2.12.0+) with an e-value cutoff of 1e-5. Contigs with more than 90% of their length aligned to sequences from microorganisms were considered contaminants and removed. Finally, contigs with sequence identity > 99% and coverage > 99% were removed. Hi-C reads were aligned to the final assembled contigs using HiCUP^45^ (v.0.6.1). Base on the Hi-C read alignments, the contigs were clustered to 28 groups using ALLHiC^46^ (v.0.9.8) with parameters ‘-e GATC -k 28’. Subsequently, to detect the collapsed regions in the assembly, Illumina paired-end reads (89 Gb) were mapped to the assembled genome using BWA-MEM^47^ with default parameters. Based on the alignments the mapping depth was calculated for each 50-kb non-overlapping window across the assembly using mosdepth^48^. Genome regions with a depth greater than 65× are identified as potential collapse regions, and contigs within these regions were added to other haplomes that lacked these regions. The final haplotype-resolved chromosome assembly results were manually corrected using JuiceBox^49^ based on Hi-C read mapping, resuling in a total of 28 haplotype-resolved chromosomes. Synteny between the assembly and the *R. chinensis* ‘Old Blush’ genome was identified using MCScanX^50^, and synteny between the assembly and the genetic map of modern rose^10^ was identified using ALLMAPS^51^ (v.0.8.12). BUSCO^8^ (v.5.4.5) and LAI^9^ (LTR_retriever; v.2.9.0) were used to evaluate the quality of the assembly.

### Repeat sequence annotation and protein-coding gene prediction

A *de novo* long terminal repeat retrotransposon (LTR-RT) library and a miniature inverted repeat transposable element (MITE) library were constructed for the assembled ‘Samantha’ genome using LTRharvest^52^ (v.1.5.10) and MITE-Hunter^53^ (v.11-2011), respectively. The ‘Samantha’ assembly was then masked using RepeatMasker^54^ (v.4.0.6) with the LTR-RT and MITE libraries. The unmasked sequences in the genome were further searched for repeat elements using RepeatModeler^55^ (v.1.0.11). All the repetitive sequences generated above were combined into one *de novo* repeat library, which was used to search the genome for repeat sequences using RepeatMasker^54^ (v.4.0.6).

The repeat-masked genome was used for protein-coding gene prediction with the MAKER pipeline^56^ (v.2.31.10), which combines evidence of *ab initio* gene prediction, transcript mapping, and protein homology to define the final gene models. SNAP^57^ (v.2006-07-28) and AUGUSTUS^58^ (v.3.3) were used for *ab initio* gene predictions. The RNA-Seq data were cleaned with Trimmomatic^59^ (version 0.39) and assembled using Trinity^60^ (v.2.4.0) with the *de novo* mode and the genome-guided mode, respectively. The assembled contigs were used as the transcript evidence. Protein sequences from *Arabidopsis*^61^, *Malus*^62^, *Fragaria*^63^, diploid rose^6^ and *Pyrus*^64^, and the UniProt database were aligned to the ‘Samantha’ genome using Spaln^65^ to provide the protein homology evidence. For gene annotation, protein sequences of the predicted genes were compared against *Arabidopsis* protein and UniProt (Swiss-Prot/TrEMBL) databases using BLAST, as well as the InterPro database using InterProScan^66^. GO annotations were obtained using Blast2GO^67^.

### Subgenome dominance analysis

A total of 62 accessions from wild original species were used to assign the four assembled haplomes to the two subgenomes. For each of the 28 assembled chromosomes, SNPs were identified in the 62 accessions using genome resequencing data with the ‘Samantha’ chromosomes as the reference and used to construct the phylogenetic trees (Supplementary Fig. 3). Haplomes phylogenetically closest to the Chinese original species were referred to as ChrA, and haplomes closest to the European original species were named ChrB. The other two haplomes closely related to Chinese and European original species, respectively, were referred to as ChrC and ChrD. OrthoVenn2^68^ was used for gene family clustering analysis to detect specific genes in each of the four haplomes.

Two haplomes, ChrA and ChrB, from each of the two subgenomes, were used for subgenome dominance analysis. Gene retention analysis was performed using the woodland strawberry genome as the reference^69^, and orthologous genes between ChrA and ChrB were identified using Liftoff^70^ (v.1.6.3). The retained homoeologous pairs were identified following the method in Schnable et al.^69^. *Ka*/*Ks* ratios of orthologous gene pairs between each of the two ‘Samantha’ subgenomes and the woodland strawberry were calculated using TBtools^71^.

RNA-Seq data from 54 different samples were used to detect the dominance of subgenomic expression, and these samples included cut flower of ‘Samantha’ under different hormone treatments. RNA-Seq reads were processed using Trimmomatic^59^ (v.0.39) to remove adapter, poly(A/T) tails, and low-quality sequences. The processed reads were then aligned to the SILVA rRNA database (release 138; https://www.arb-silva.de/) to remove rRNA contaminations. Cleaned reads were aligned to the two subgenomes of ‘Samantha’ using HISAT2^72^ (v.2.2.1) with default parameters. Only uniquely mapped reads were retained, and the read counts in 1-Mb non-overlapping windows were calculated using SAMtools^73^ (v.1.17).

Syntenic tetrads among the four haplomes (one from each of the four haplomes) were identified and used to analyze expression bias. Kallisto^74^ (v.0.48.0) was used to calculate the expression of genes with default parameters. For each transcriptome, only those quadruple homologous genes with at least one expressed were included in the analysis. The definition of expression bias was referred to the study of polyploid wheat^75^.

### Variant calling and annotation

The Illumina paired-end reads from each rose accession were processed to remove adaptor and low-quality sequences using Trimmomatic^59^ (v.0.39). The cleaned reads were aligned to subgenomes ChrA and ChrB, respectively, using BWA MEM^47^ (v.0.7.17) with default parameters. Picard (v2.7.1; http://broadinstitute.github.io/picard/) was used to mark duplicated alignments. The HaplotypeCaller function in GATK^76^ (v.4.2.6) was then used to generate a GVCF file for each of the 233 rose accessions. All GVCF files were combined for variant calling using the function GenotypeGVCFs in GATK^76^. Hard filter was applied to the identified raw SNPs using GATK with parameters ‘QD < 2.0 || FS > 60.0 || MQ < 40.0’. SNPs were further filtered to keep those that were biallelic in the population, had an average mapping rate ≤ 1.5 times of the average genome-wide mapping rate. SNP annotation was performed using the package ANNOVAR^77^ (v.2015-12-14).

### Phylogenetic and population genomic analysis

We selected a total of 14,060,480 SNPs with minor allele frequency (MAF) ≥ 5% and missing data rate ≤ 10% for phylogenetic and population structure analyses. A neighbor-joining phylogenetic tree was constructed using PHYLIP^78^ (v.3.5). PCA was performed using PLINK^79^ (v.1.90b6.10). Population structure was investigated using ADMIXTURE^80^ (v.1.3.0) with default parameters. To determine the most likely number of ancestral kinships (*K*) in the rose population, *K* values were set from 2 to 10. The statistic ‘cross-validation error (CV error)’, which indicates the change in likelihood of different numbers of clusters, was calculated, and the cluster number with the lowest CV error, which indicates the most likely number of clusters in the population, was obtained.

Nucleotide diversity (π) were calculated in 500-kb non-overlapping windows across the ‘Samantha’ genome with the final set of 14,060,480 SNPs using vcftools^81^ (v.0.1.16). Fixation index (*F*_ST_) values were calculated between two groups (tetraploid intermediate old cultivars and tetraploid modern cultivars) using vcftools^81^ (v.0.1.16).

LD decay patterns were calculated using PopLDdecay^82^ (v.3.42) with the default parameters for the two groups (tetraploid intermediate old cultivars and tetraploid modern cultivars). Correlation coefficient (*r*^2^) of genotypes was used as a measure of the LD level. SNPs within each group were extracted for the analysis. SNPs within 2-kb sliding windows were used to estimate average *r*^2^ at various physical distance classes, and the LD decay was plotted as a function of the derived average *r*^2^ and the physical distances along the genome.

### Selective sweep identification

We identified the selection signals across the two subgenomes based on the π ratios and *F*_ST_ values. Selective sweep screening was performed by comparing tetraploid intermediate old cultivars (Int_4; n = 45) vs. tetraploid modern cultivars (Hyb_4; n = 32). A 50-kb sliding window with 5-kb step approach was applied to quantify *F*_ST_ and π ratios. Regions with the top 5% of both *F*_ST_ values and π ratios were considered as candidate selective sweeps.

To infer potential sources of interesting selective sweeps in tetraploid modern cultivars, 55 accessions from original species with a single genetic component greater than 95% at *K*=9 in population structure analysis, were used for SNP genotyping analysis. SNPs in each selective sweep were used to construct the ML phylogenetic tree with IQ-TREE^83^ (v.1.6.8). The genotypes of SNPs in each selective sweep region were presented through a heatmap, which was generated using the R package pheatmap (v.1.0.12).

### Genomic composition analysis

Six potential original species (*R. chinensis* ‘Old Blush’, *R. odorata* var. *gigantea*, *R. gallica*, *R. wichuraiana*, *R. rugosa* and *R. moschata*) of the ‘Samantha’ genome identified by ADMIXTURE^80^ were used for genetic organization analysis. The original attribution of chromosomes was inferred through two approaches: (1) Equal amount of Illumina short read data from each of the six species were aligned to the four haplomes of the ‘Samantha’ genome. Within each of the 500-kb non-overlapping windows, if the highest sequencing coverage depth among the six species was 1.5 times higher than the second highest, this window was considered to be attributed to the species with the highest sequencing coverage depth. (2) High-quality SNPs were called based on the alignments, and the potential original attribution of the 500-kb non-overlapping windows was inferred by counting the proportion of homozygous variants between each species and the four haplomes of ‘Samantha’. For each specific genomic region of ‘Samantha’, species with the lowest proportion of homozygous variants was considered to be the original contributor of this region. Only results obtained by the two approaches that were consistent were used to determine the final contributing species of each region across the ‘Samantha’ chromosomes.

### Data availability

Raw reads generated in this study have been deposited in NCBI BioProject database under accession numbers PRJNA957905, PRJNA704782, and PRJNA964585. Other RNA-seq data used in this study were downloaded from the NCBI Bioproject database under the accession numbers PRJNA486271^84^ and PRJNA522664^85^. The sequences and annotations of the ‘Samantha’ genome assembly are available at Figshare (https://doi.org/10.6084/m9.figshare.22774097).

## Acknowledgements

We thank Dr. Tao Lin (China Agricultural University) and Dr. Qiang Gao (BGI Tech., Beijing) for helpful discussions, Dr. Haibo Xin and Dr. Yanhua Bu (Beijing Institute of Landscape Architecture) for providing computing resources and *Rosa* materials, and the Center of Agricultural Biotechnology of Beijing Academy of Agriculture and Forestry Sciences for the help of flow cytometry experiments. This work was supported by funds from the 111 Project of the Ministry of Education (B17043 to J.G.), the Construction of Beijing Science and Technology Innovation and Service Capacity in Top Subjects (CEFF-PXM2019_014207_000032 to J.G.), National Key Research and Development Program of China (Grant no. 2018YFD1000400 to J.G.), General Project of Shenzhen Science and Technology and Innovation Commission (21K270360620 to Y.Li) and National Natural Science Foundation of China (Grant no. 31572162 to Y.Li, 31772344 and 31972444 to Z.Z, and 31522049 and 31872148 to N.M.).

## Author contributions

J.G., N.M. and Z.F. designed and coordinated the project. Y. Liu, W.W., H.R. and S.S. performed DNA extraction. Y. Liu performed the flow cytometry analysis. Y.Y., L.L., S.D., Y. Zhu, Y.C., H. Zhou., H. Zhang., J.C, and K.T. contributed *Rosa* materials. Z.Z., Y. Liu., Y. Li., T.Y., Q.P., X.S., Y.J. and X.Z. coordinated sample collection and sequence data generation. D.G., L.C., S.W., S.S., H.S., J.W. and Y.Z. integrated the genome assembly and annotation. H.S., T.Y. performed subgenome dominance and selective sweep analysis. H.S., J.W., T.Y. and Y.Z. conducted phylogenetic and population genomic analysis. T.Y., Y. Liu, J.W., S.W., Z.Z., Y.L., Z.F., N.M., and J.G. wrote and revised the manuscript.

