## Supplementary Figure for "Haplotype-resolve genome assembly and resequencing provide insights into the origin and domestication of modern rose"

### Supplementary Note 1. Old rose cultivars in Europe

#### 1.1 European old cultivars

Before the introduction of Chinese species/cultivars, roses cultivated from wild species native to Europe/Middle East can be classified into the following groups:

| Group | Description |
|---|---|
| Gallica | Once-flowering. Bred from wild species *R. gallica*. |
| Damask | Mostly once-flowering, except 'Autumn Damask' that can sporadically flower in autumn. Known for strong fragrance. Recorded to be hybrid of *R. gallica* and other uncertain species possibly including *R. phoenicia*, *R. moschata*, *R. canina*, *R. fedtschenkoana* and/or *R. damascena*. |
| Alba | Once-flowering. Known for their light (white or pink) flower colors and elegant fragrance. Believed to be hybrids of Damask roses and wild species *R. canina*. |
| Centifolia | Mostly once-flowering. Believed to be hybrids of Damask (or *R. gallica*) and Alba roses. Often having more than 100 petals, and thus referred to as 'centifolia' and 'cabbage rose'. |
| Moss | Mostly once-flowering. Known for their mossy glandular growth on sepals, peduncle, and stems and wherefrom oleoresin secreted. Believed to be sports of Centifolia. |
| Hybrid foetida | Hybrid of wild species *R. foetida*. |

#### 1.2 Intermediate old cultivars

After the introduction of Chinese species/cultivars to Europe, they were extensively used in rose hybridization, especially for backcrosses to obtain the continuous-flowering trait. These intermediate cultivars can be classified into the following groups:

| Group | Description |
|---|---|
| Bourbon | First repeat-flowering cultivar group after old Chinese cultivars. Found in Ile de Bourbon. Maybe hybrids of 'Autumn Damask' and old Chinese cultivars. |
| Portland | Continuous-flowering. Maybe hybrids of 'Autumn Damask' and 'Slater's Crimson China'. |
| Noisette | Continuous-flowering. First Noisette cultivar 'Champney's Pink Cluster' is a hybrid of *R. moschata* and *R. chinensis* 'Old Blush'. Known for blooms in fragrant clusters. |
| Hybrid Multiflora | Once-flowering. Bred from Asian wild species *R. multiflora*. Known for blooms in large clusters. |
| Tea | Continuous-flowering. Bred from *R. odorata* cv. 'Hume's Blush Tea-scented China' and/or 'Park's Tea-scented China'. Known for its flower shape. The forerunner of modern Hybrid Teas. |

| | |
|---|---|
| Hybrid Perpetual | Continuous-flowering. A further step from Bourbon to modern Hybrid Teas. Hybrids of Portland, Noisette, Bourbon, and Tea roses. Known for their steady continuous-flowering and diverse colors. |

### Supplementary Note 2. Sections within subgenus *Rosa*

According to Flora of China[1], there are nine sections within subgenus *Rosa* distributed in China: Chinenses, Synstylae, Cinnamomeae, Pimpinellifoliae, Microphyllae, Banksianae, Bracteatae, Laevigatae and Rosa. In addition, we included two accessions from the section Caninae.

| **Section** | **Description** |
|---|---|
| Chinenses | Include three species, *R. chinensis*, *R. odorata*, and *R. lucidissima*. Include famous old cultivars R. *chinensis* cv. 'Old Blush', *R. chinensis* cv. 'Slater's Crimson China', *R. odorata* cv. 'Hume's Blush Tea-scented China', and *R. odorata* cv. 'Park's Yellow Tea-scented China' that were introduced into Europe and greatly flourished rose breeding in Europe. |
| Synstylae | Mainly distributed in Asia. Include 18 species distributed in China, among which *R. multiflora* and *R. wichuraiana* are the most commonly used in rose breeding. The most obvious traits are their inflorescence and the combination of style, giving them the name of the section. |
| Cinnamomeae | Widely distributed in Asia, Europe and America. Include 31 species distributed in China, among which *R. fedtschenkoana* and *R. rugosa* were recorded to be involved in rose breeding. |
| Pimpinellifoliae | Mainly distributed in Asia and Europe. Include 18 species distributed in China. Include four-petal (e.g., *R. sericea*) or five-petal (e.g., *R. foetida*) species. |
| Microphyllae | Include three species, *R. roxburghii*, *R. kweichowensis*, and *R. praelucens*, distributed in China. *R. praelucens*, endemic to Zhongdian Plateau, Yunnan Province, is the naturally occurring decaploid grown on the highest altitude among all roses. |
| Banksianae | Include two species, *R. banksiae* and *R. cymosa*, distributed in China. |
| Bracteatae | Include two species in Asia, of which *R. bracteata* is distributed in China. |
| Laevigatae | Include only one species, *R. laevigata*, distributed in China. |
| Rosa | Originally distributed in Europe and West Asia and then introduced into China. Species including *R. gallica*, *R. damascena*, *R. centifolia*, and *R. alba* are best known for their strong growth vigor, winter-hardy, and high bushes. |
| Caninae | Include about 50 species in Europe. Characterized by the Caninae type meiosis with unbalanced heterogamous fully sexual reproduction. |

### Supplementary Note 3. Classification of modern rose cultivars

Modern rose cultivars can be classified into following groups:

| **Group** | **Description** |
|---|---|

| | |
|---|---|
| Hybrid Tea | Accepted as the most popular group. Best known for its shapely bloom (high-centered with central cone formed and petal edge could get reflexed at later opening stages) and single long stem. Often used as cut roses. Large-flower Hybrid Tea can also be called Grandiflora. |
| Floribunda | Bears its flowers in cluster/truss. Unrivalled for its large-quantity, long-lasting flowering, though flower shape is inferior to that of Hybrid Tea. |
| Climber & Rambler | Named after their growth habit of long-arching climbing canes. With a wide range of flower shapes and colors. |
| Miniature | Average plant height is around 35-75 cm. Flower form and foliage are indeed miniature versions of both Hybrid Tea and Floribundas. |
| Shrub | Bushy roses that do not fit into above classes. |

### Supplementary Figures

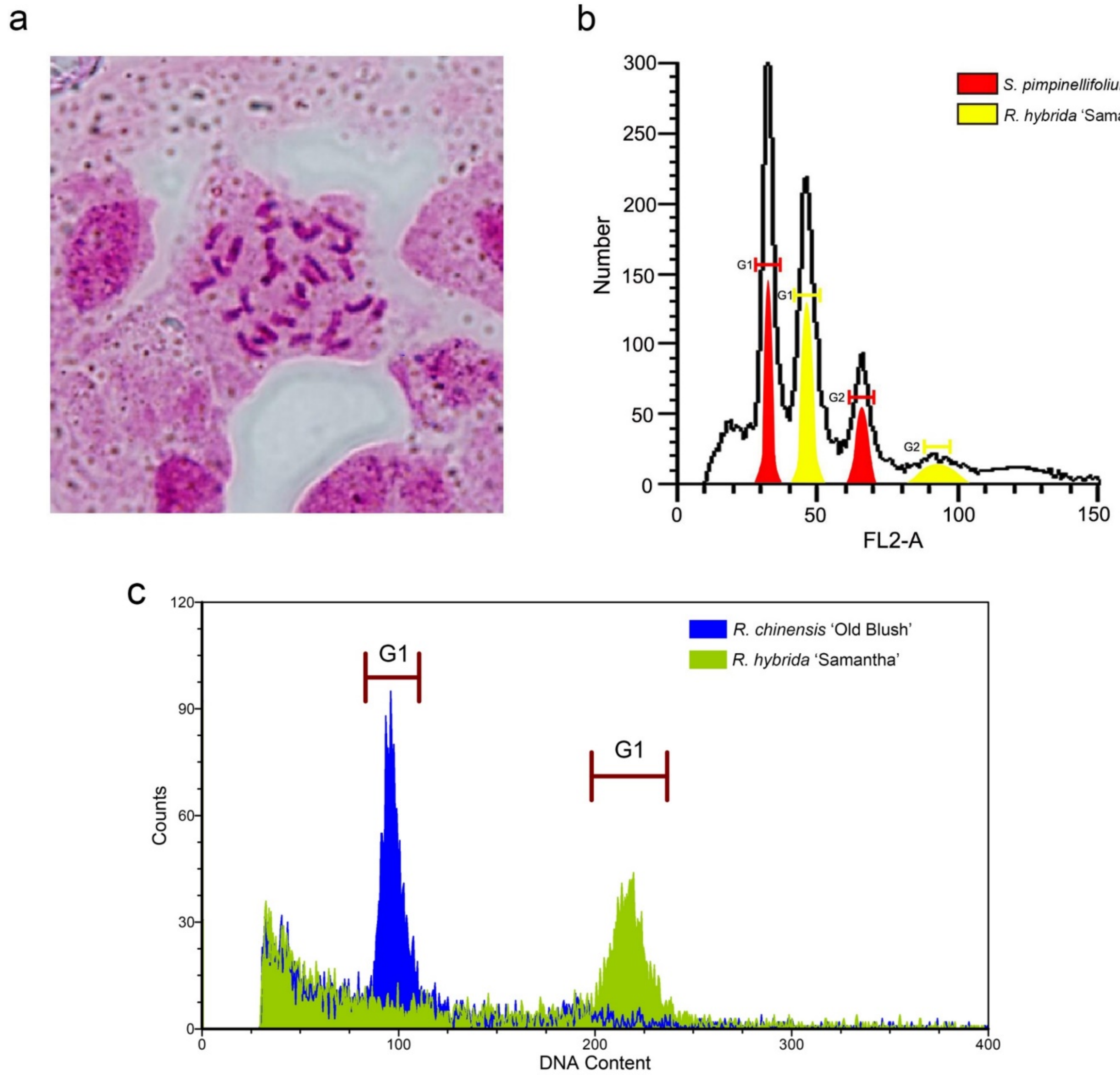


**Supplementary Figure 1. Chromosome number and genome size estimation of *R. hybrida* 'Samantha'. a**, Chromosome counting of 'Samantha' shown in a representative root cell picture. **b**,**c**, C-value estimation of 'Samantha' by flow cytometry using *Solanum pimpinellifolium* (**b**) and *R. chinensis* 'Old Blush' (**c**) as the internal reference standards. G1 peak X-values are listed in Supplementary Table 1.

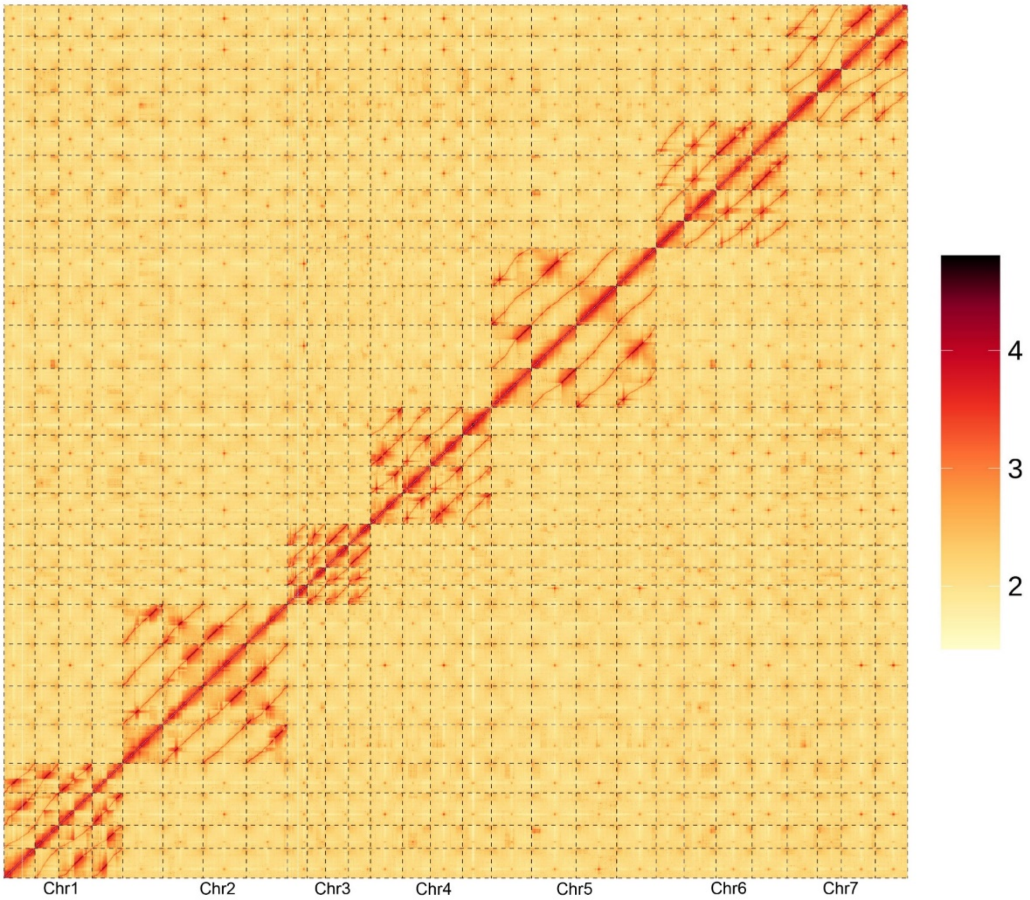


**Supplementary Figure 2.** Hi-C heatmap of the assembled *R. hybrida* 'Samantha' genome.

### Chromosome 1

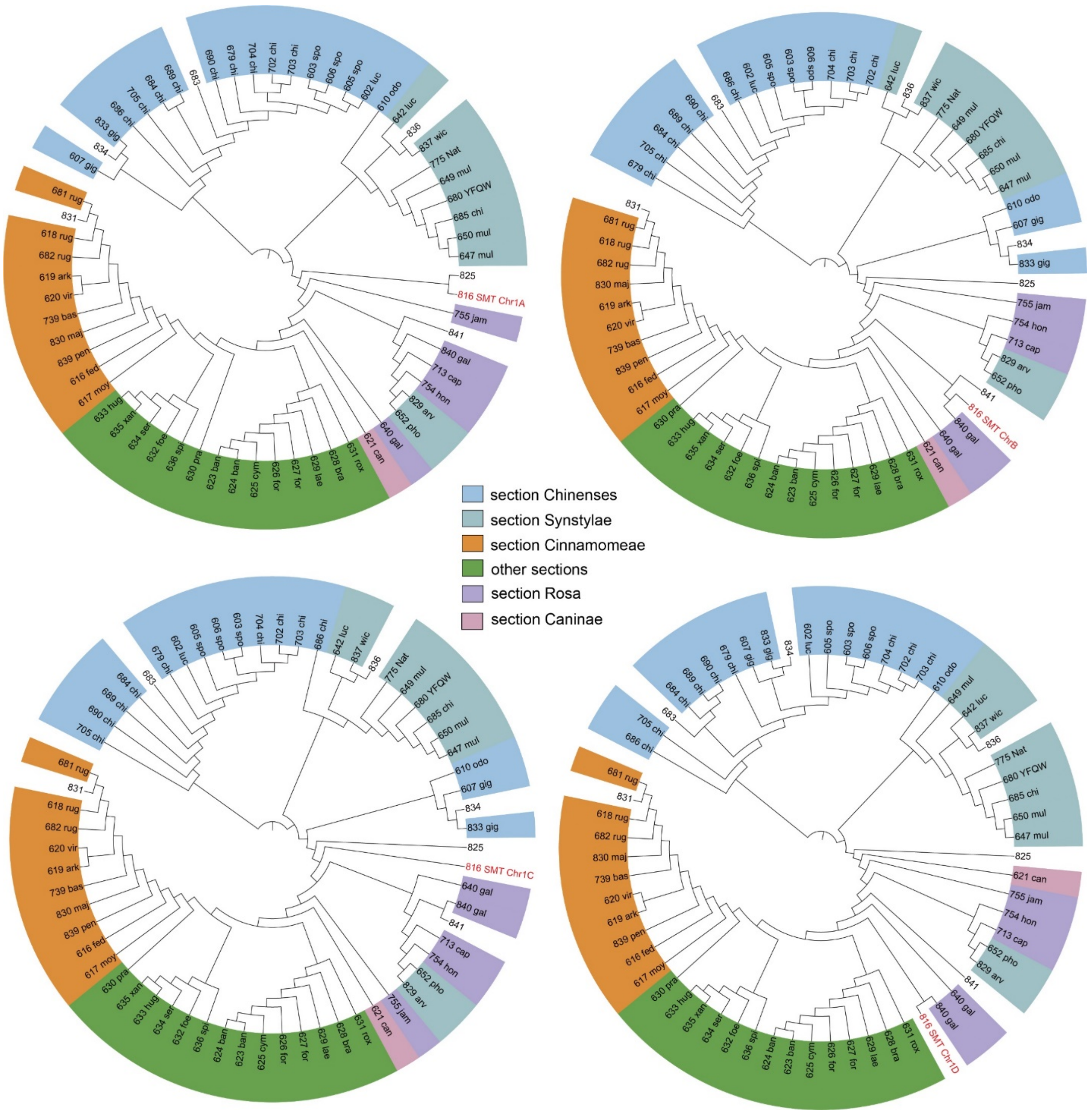


**Supplementary Figure 3.** Maximum likelihood phylogenetic trees of each of the 28 chromosomes and the original species. Genome sequencing data of 62 accession from original wild species were used for phylogenetic tree construction. ‘Samantha’ chromosomes are shown in red.

### Chromosome 2

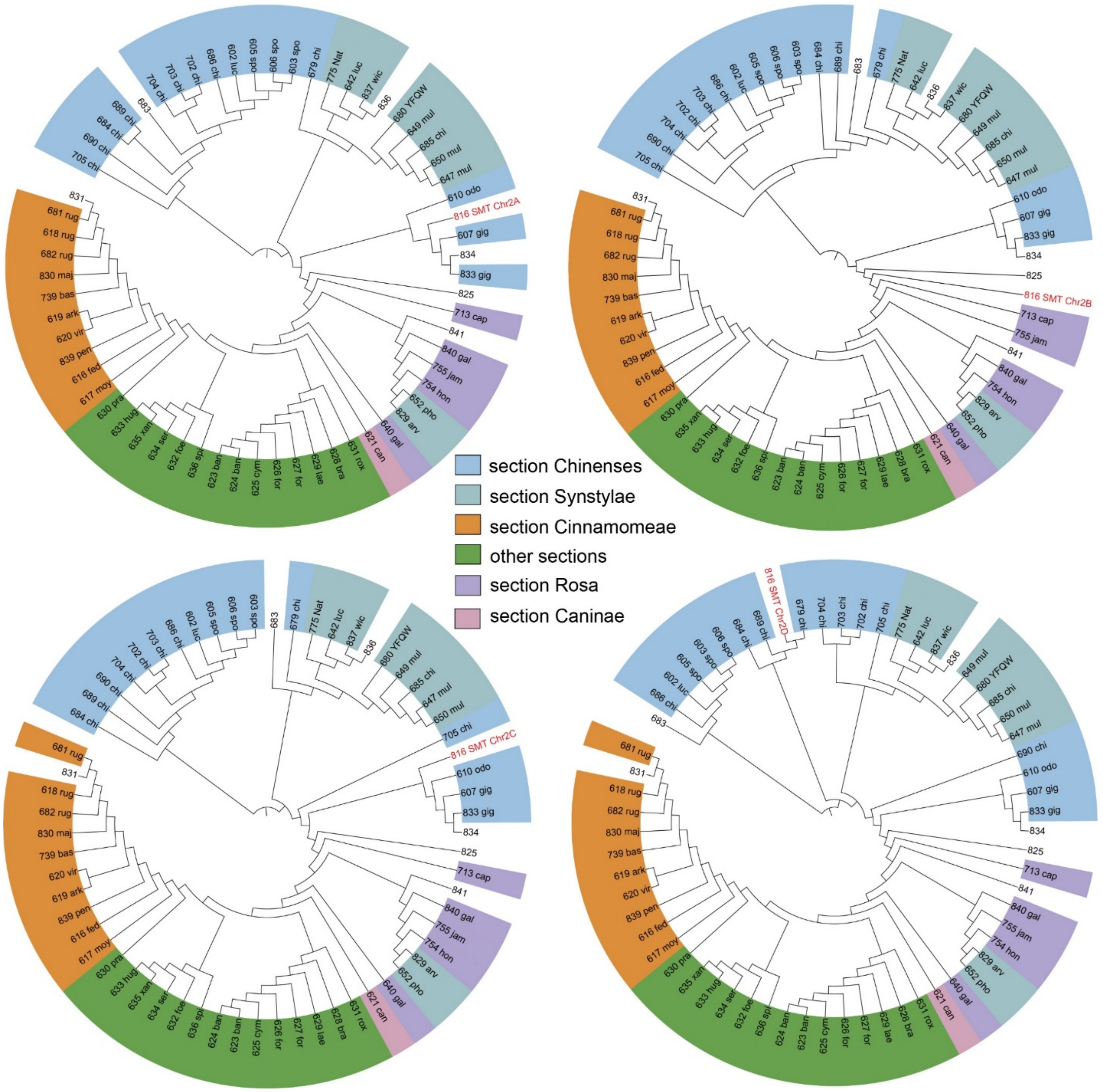


**Supplementary Figure 3.** Continued.

### Chromosome 3

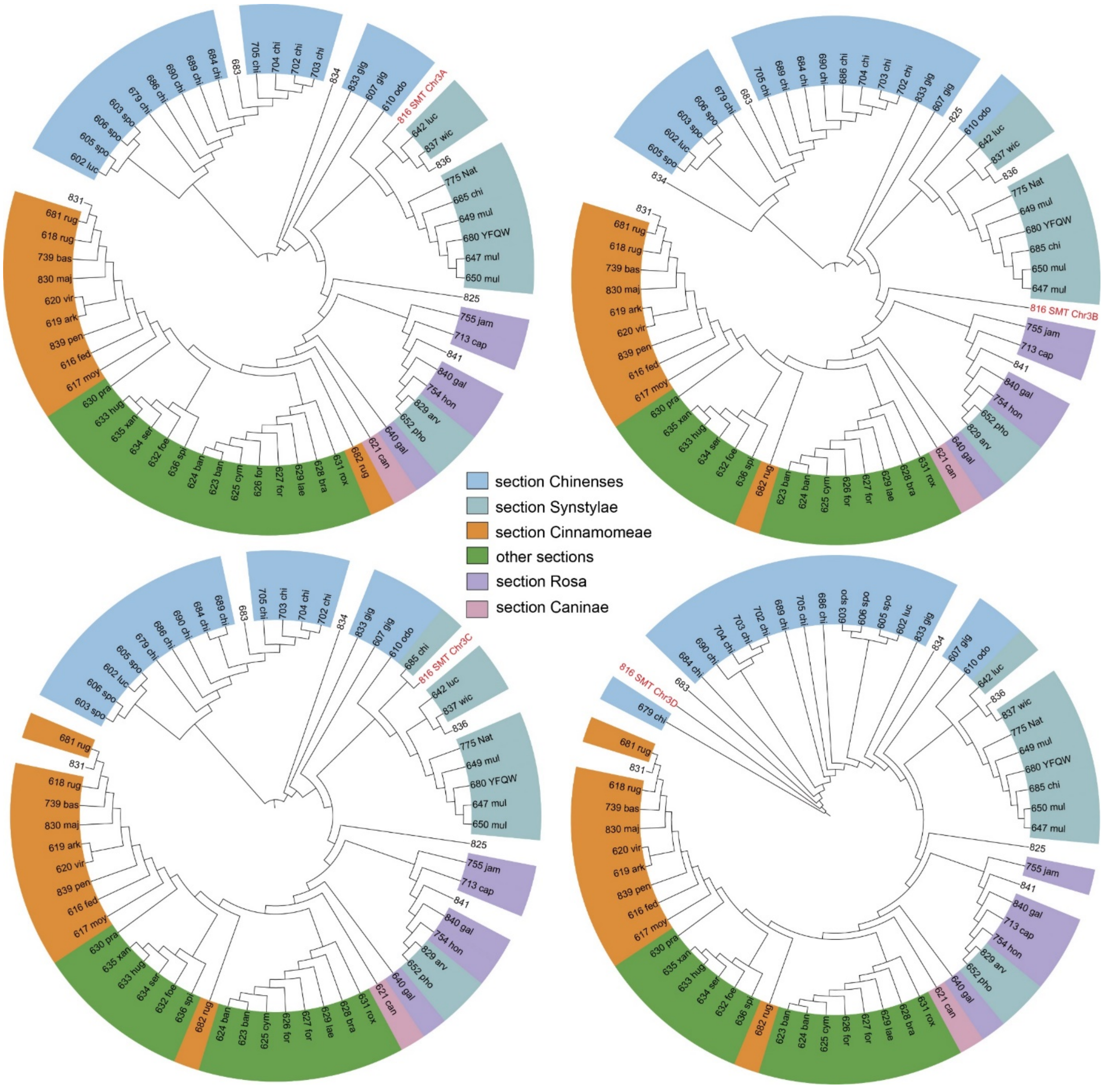


**Supplementary Figure 3.** Continued.

### Chromosome 4

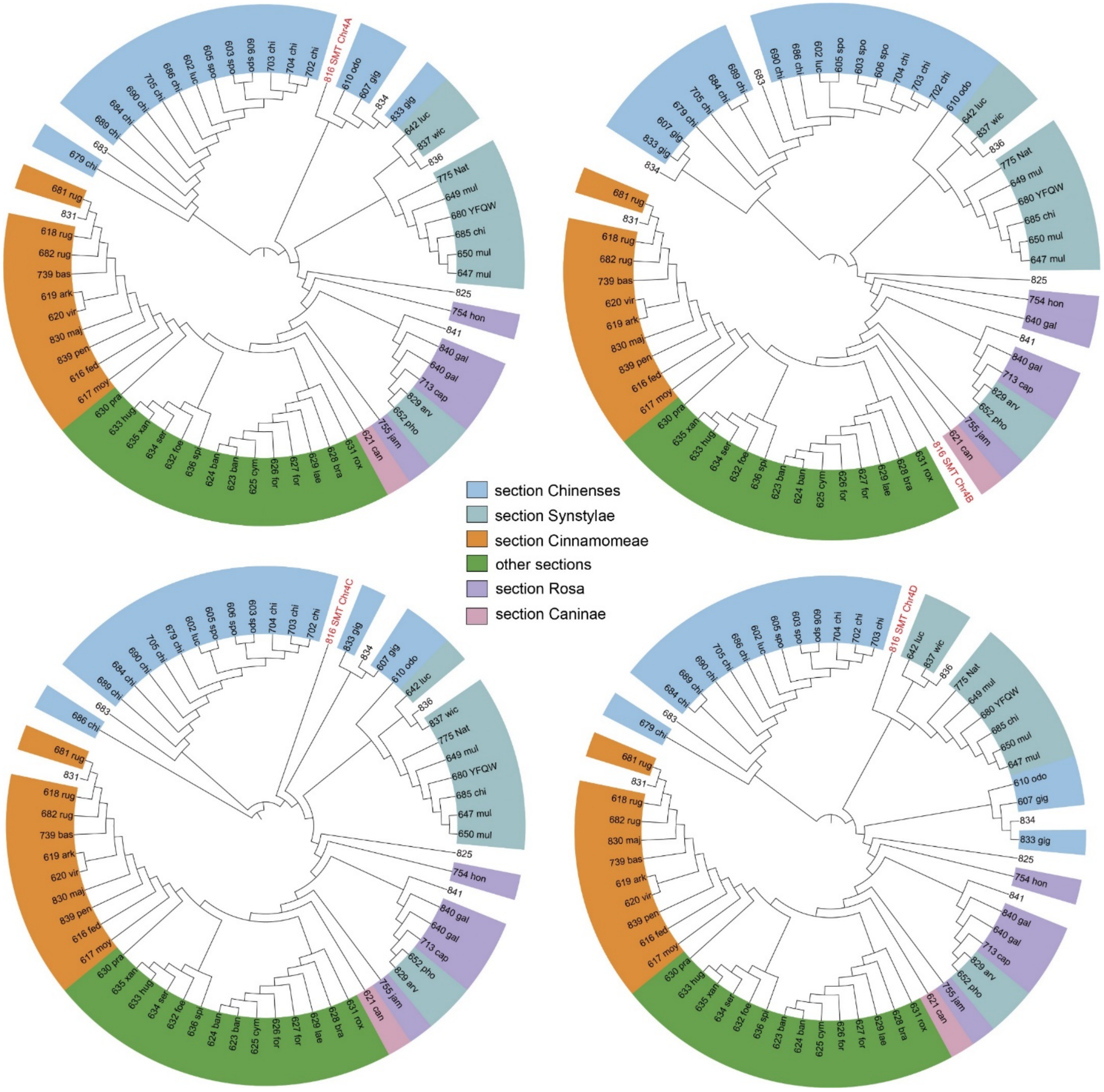


**Supplementary Figure 3.** Continued.

### Chromosome 5

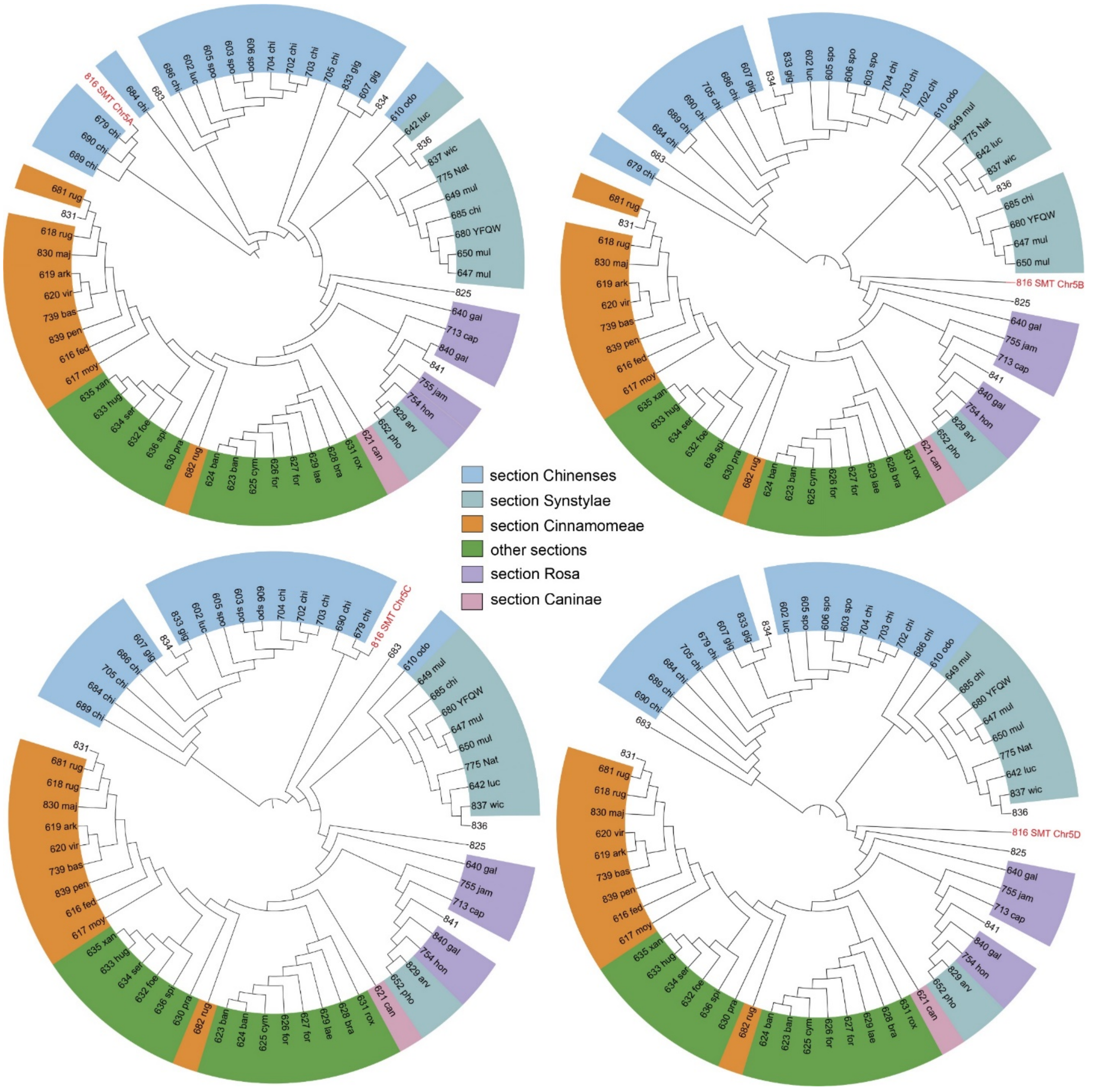


**Supplementary Figure 3.** Continued.

**Chromosome 6**

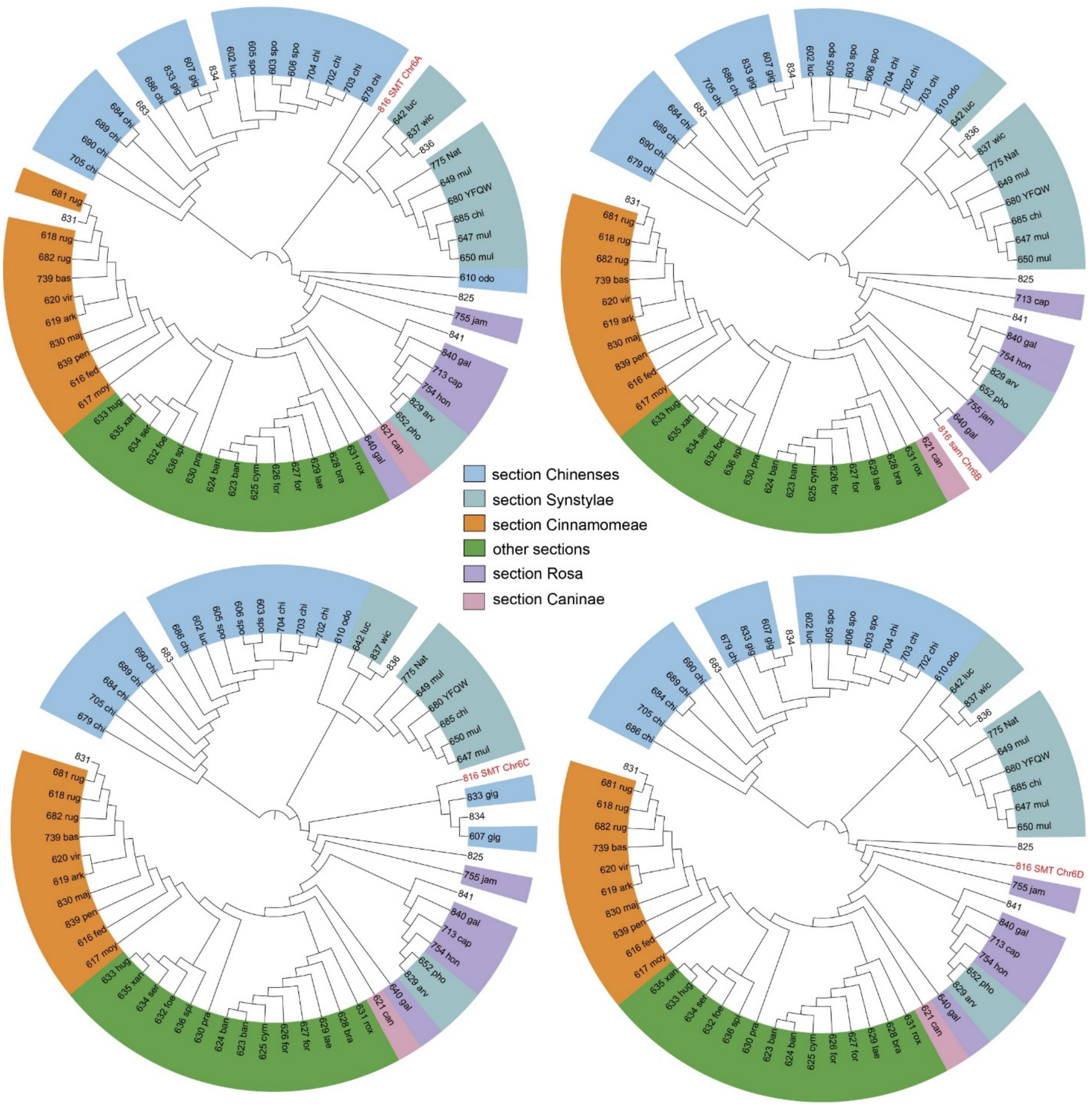


**Supplementary Figure 3.** Continued.

**Chromosome7**

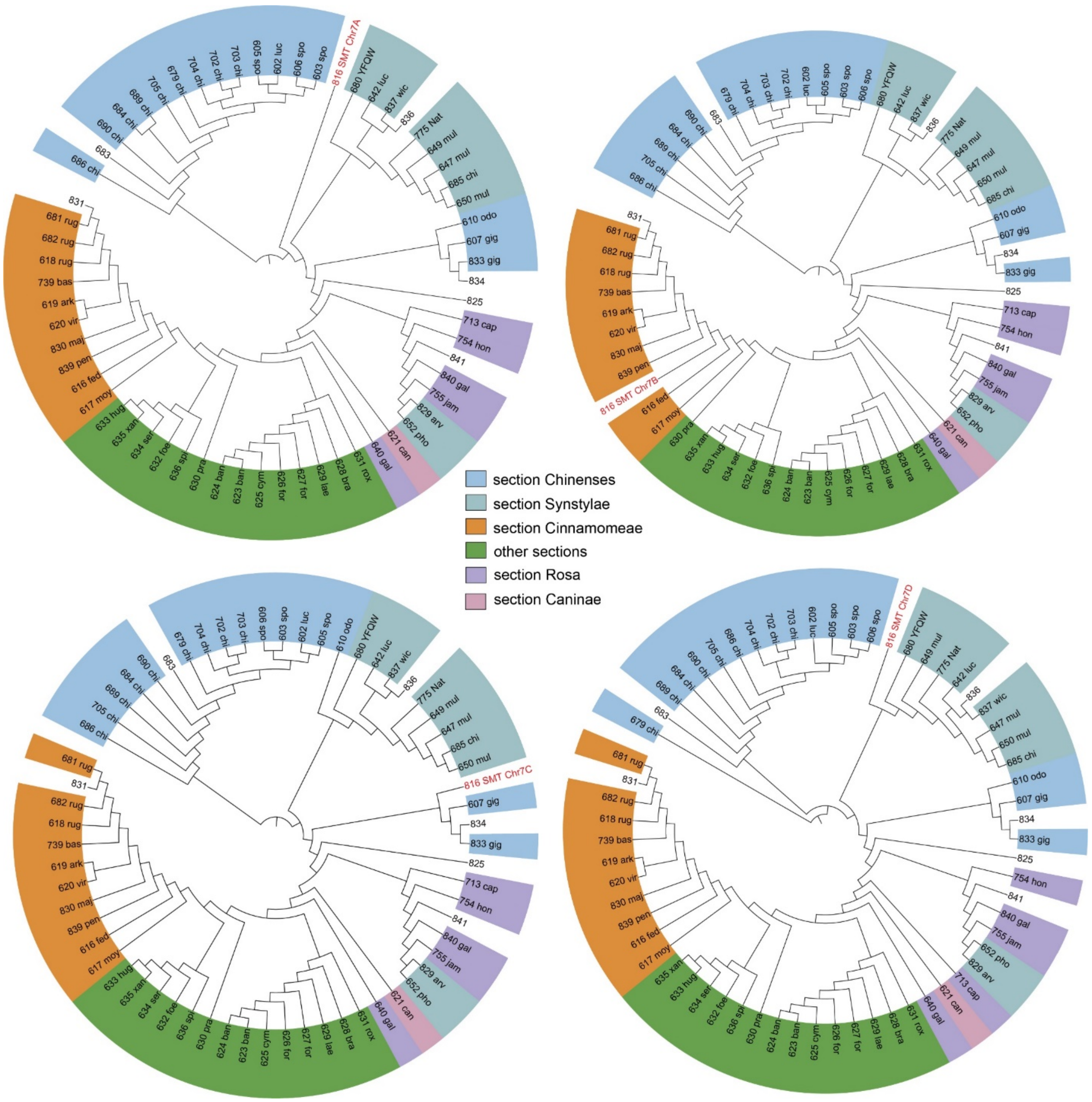


**Supplementary Figure 3.** Continued.

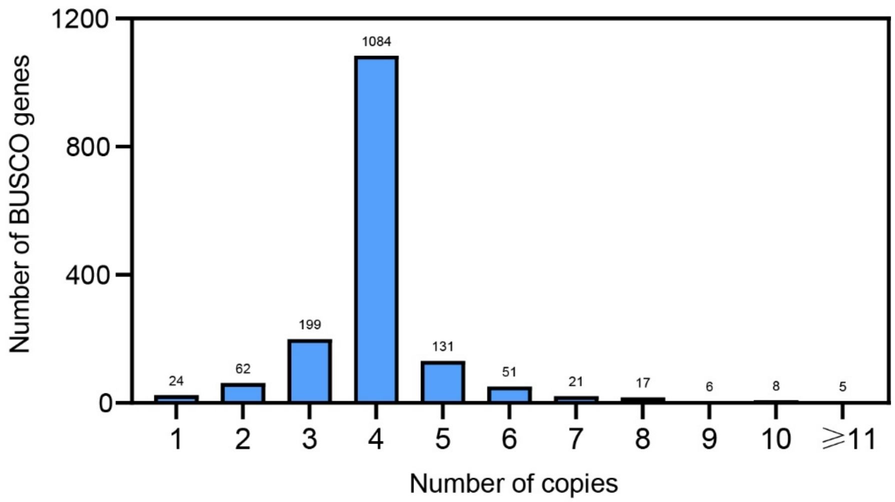


**Supplementary Figure 4.** Distribution of the number of core BUSCO genes completely captured in the ‘Samantha’ genome assembly.

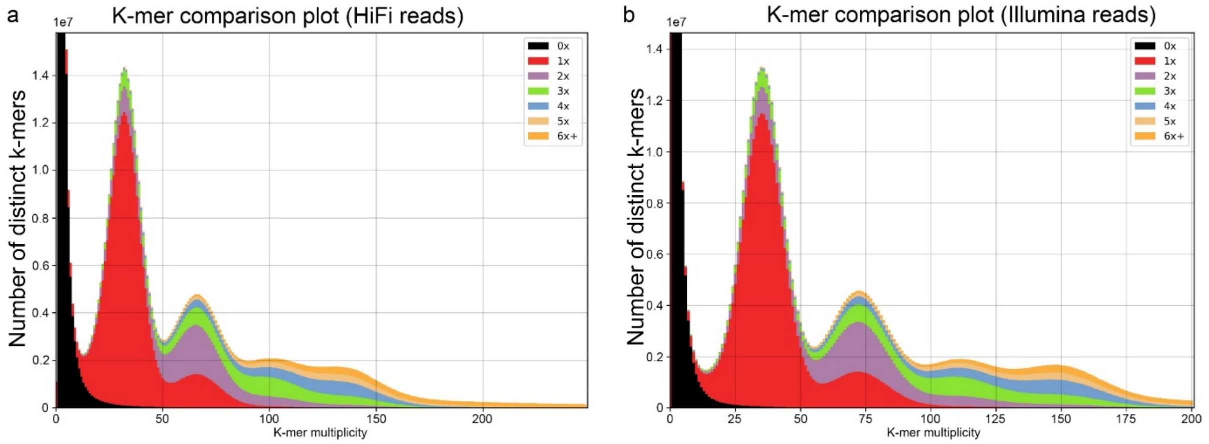


**Supplementary Figure 5.** *K*-mer spectrum analysis. Comparison 27-mer spectra between HiFi reads and the 'Samantha' assembly (**a**), and between Illumina reads and the 'Samantha' assembly (**b**). Different colors indicate different copies of the 27-mers in the genome assembly.

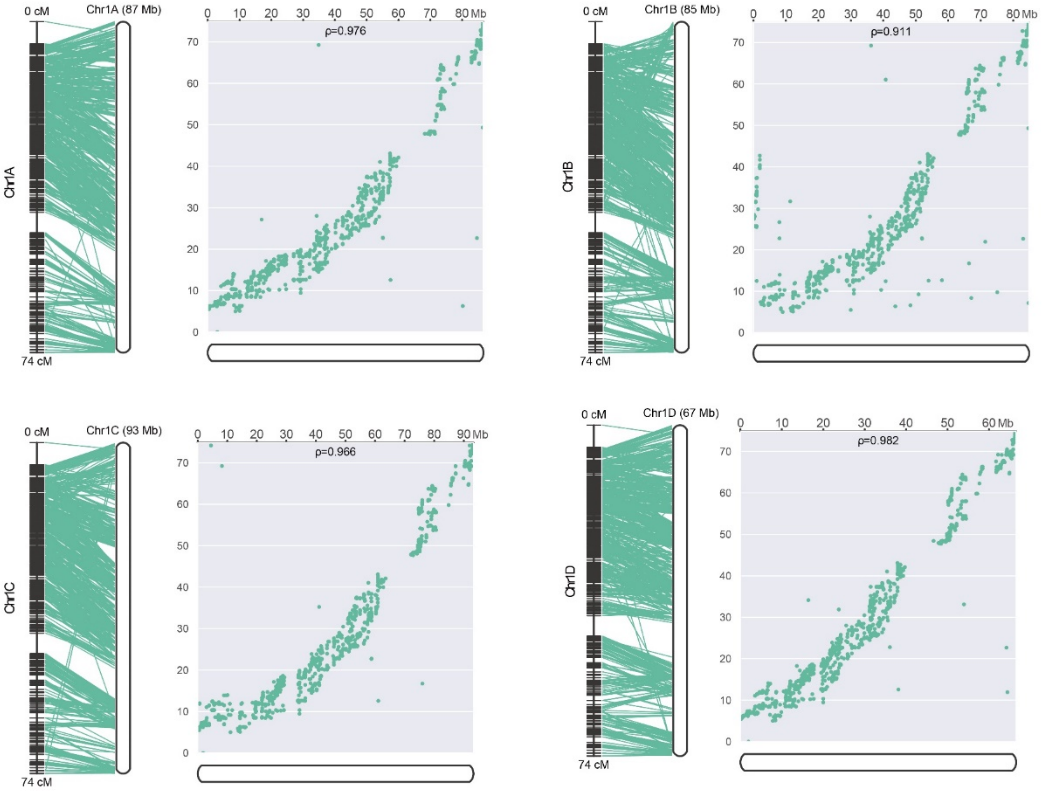


**Supplementary Figure 6.** Collinearity of the assembled 'Samantha' chromosomes with the genetic map. The left panels show the assembled chromosomes and corresponding linkage groups. The right panels show scatterplots of marker distance (cM) and physical distance (Mb). Rho (ρ) in scatterplots represents the Pearson's correlation coefficient between marker distance and physical distance.

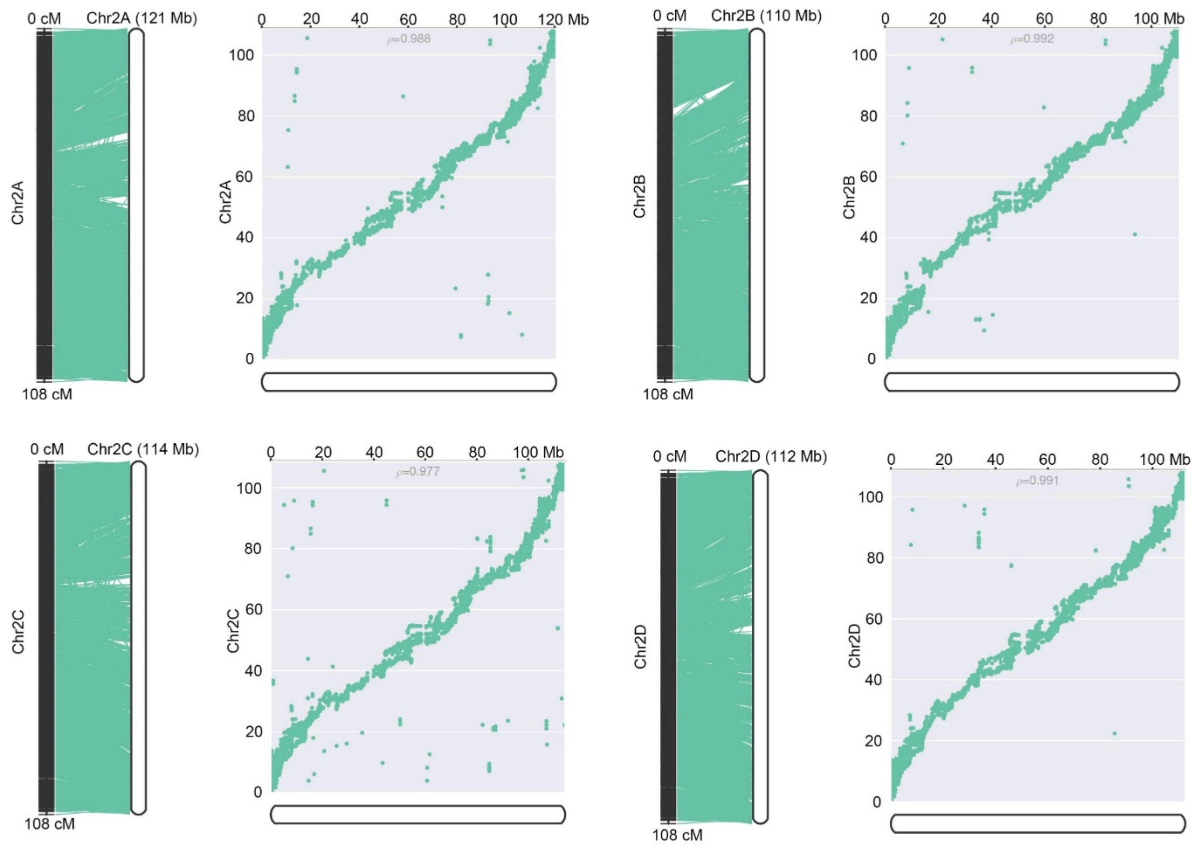


**Supplementary Figure 6.** Continued.

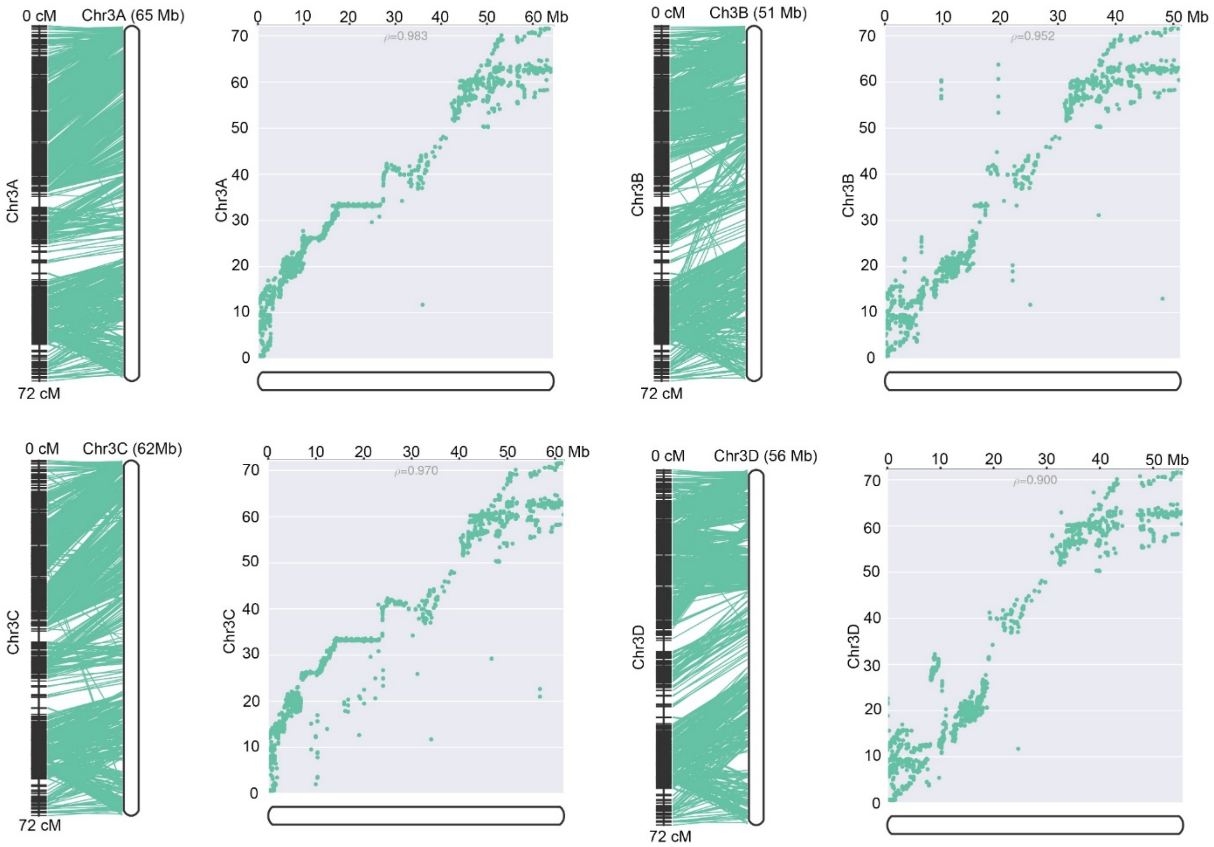


**Supplementary Figure 6.** Continued.

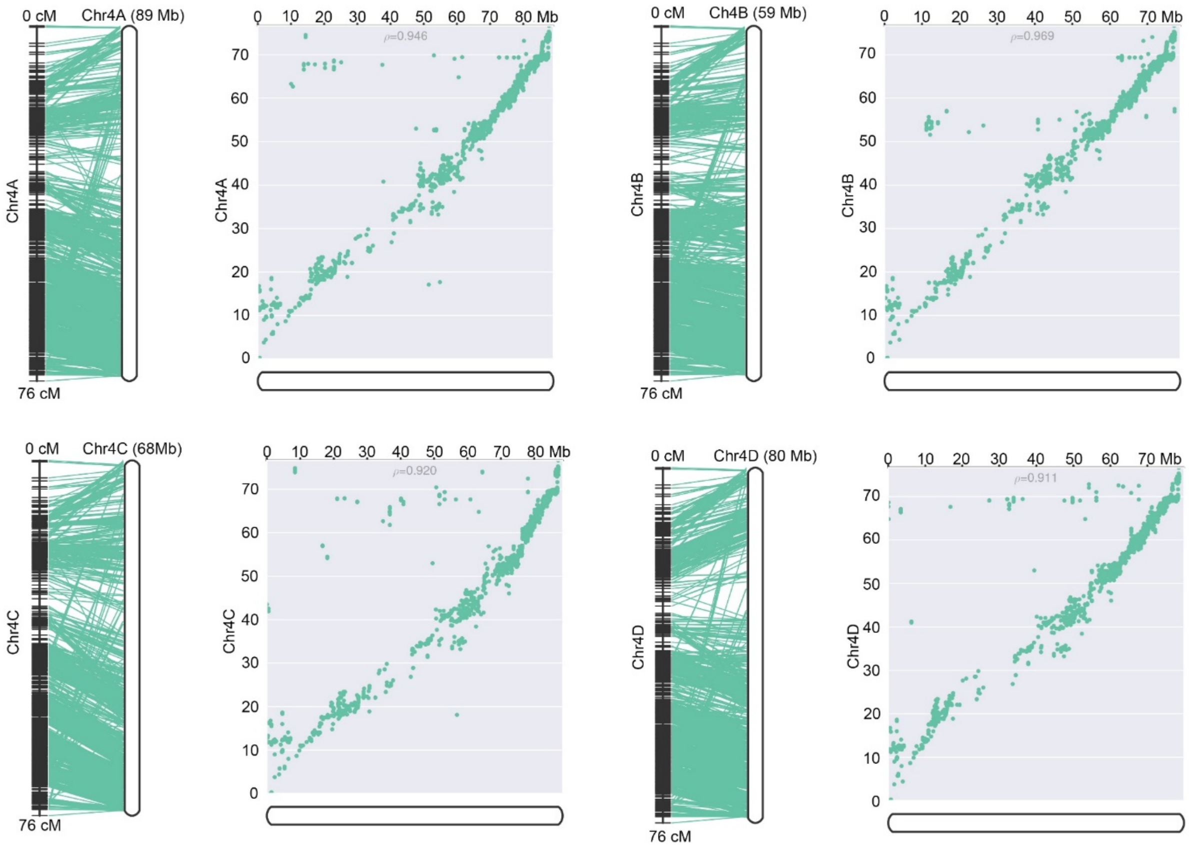


**Supplementary Figure 6.** Continued.

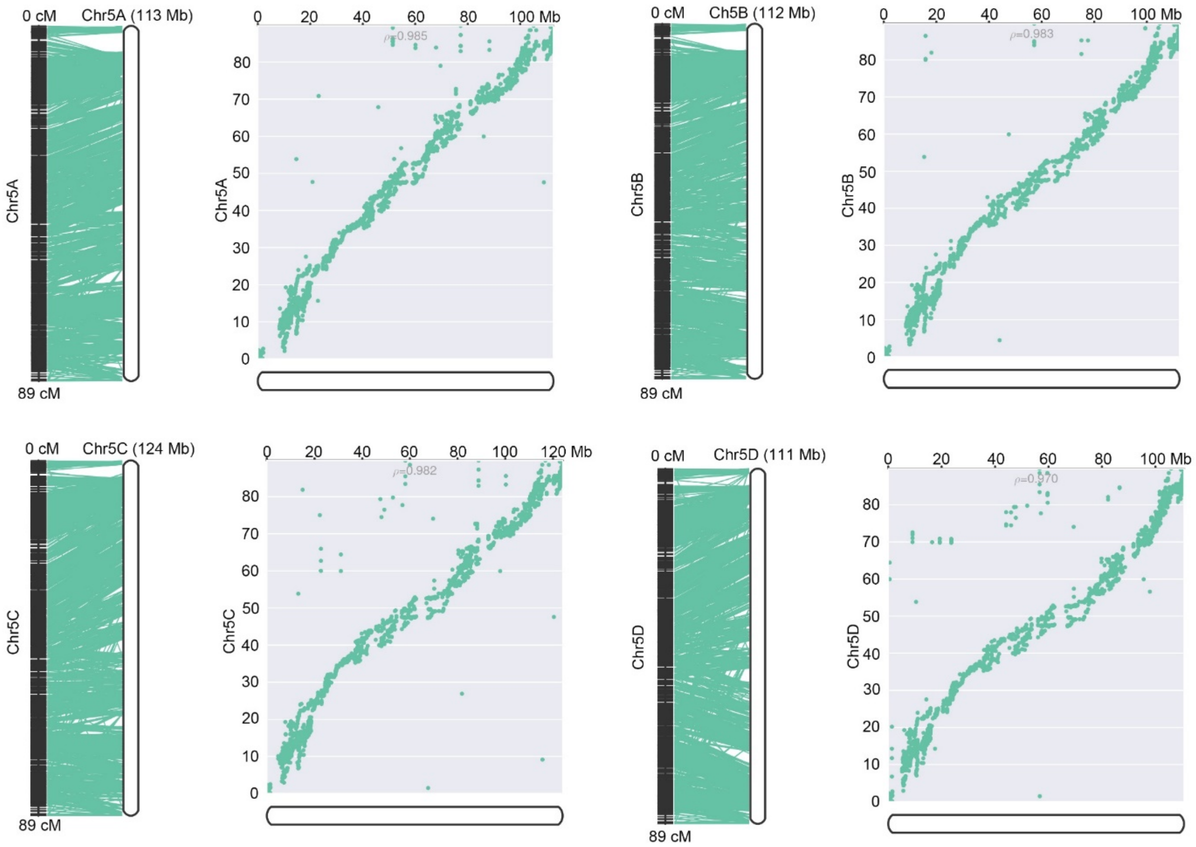


**Supplementary Figure 6.** Continued.

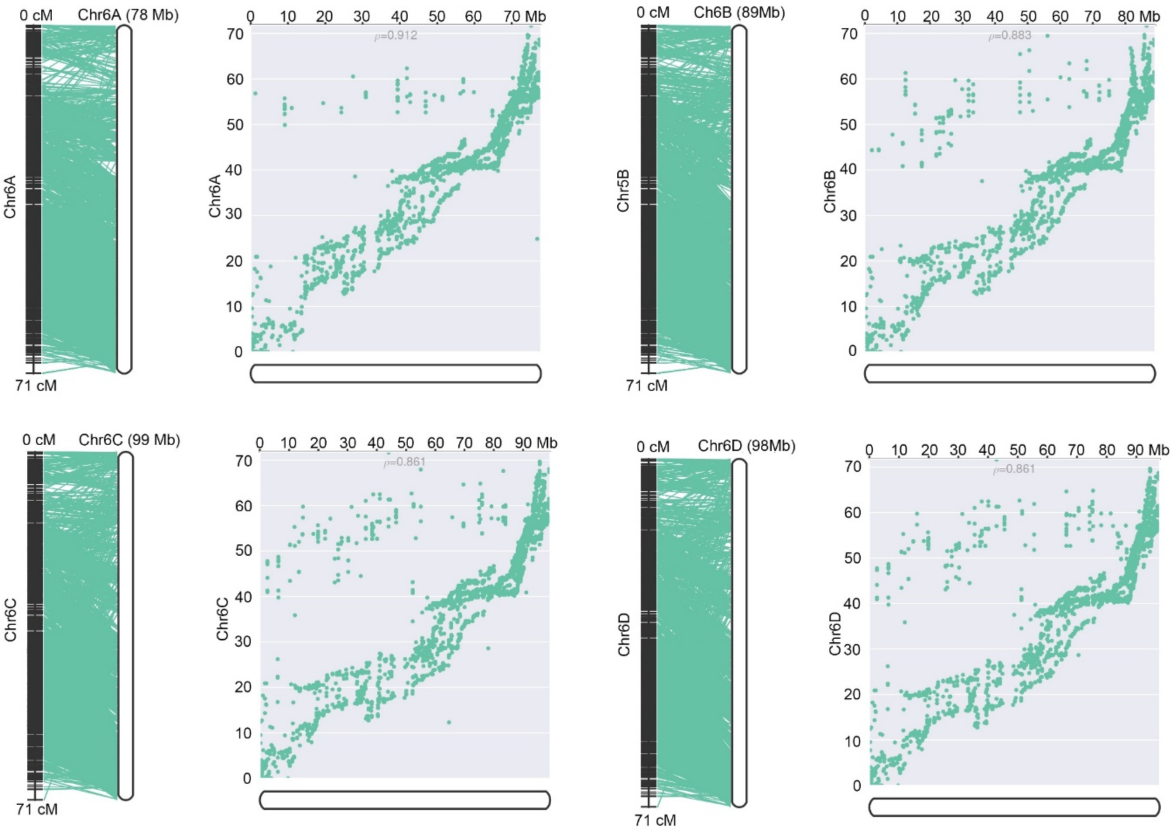


**Supplementary Figure 6.** Continued.

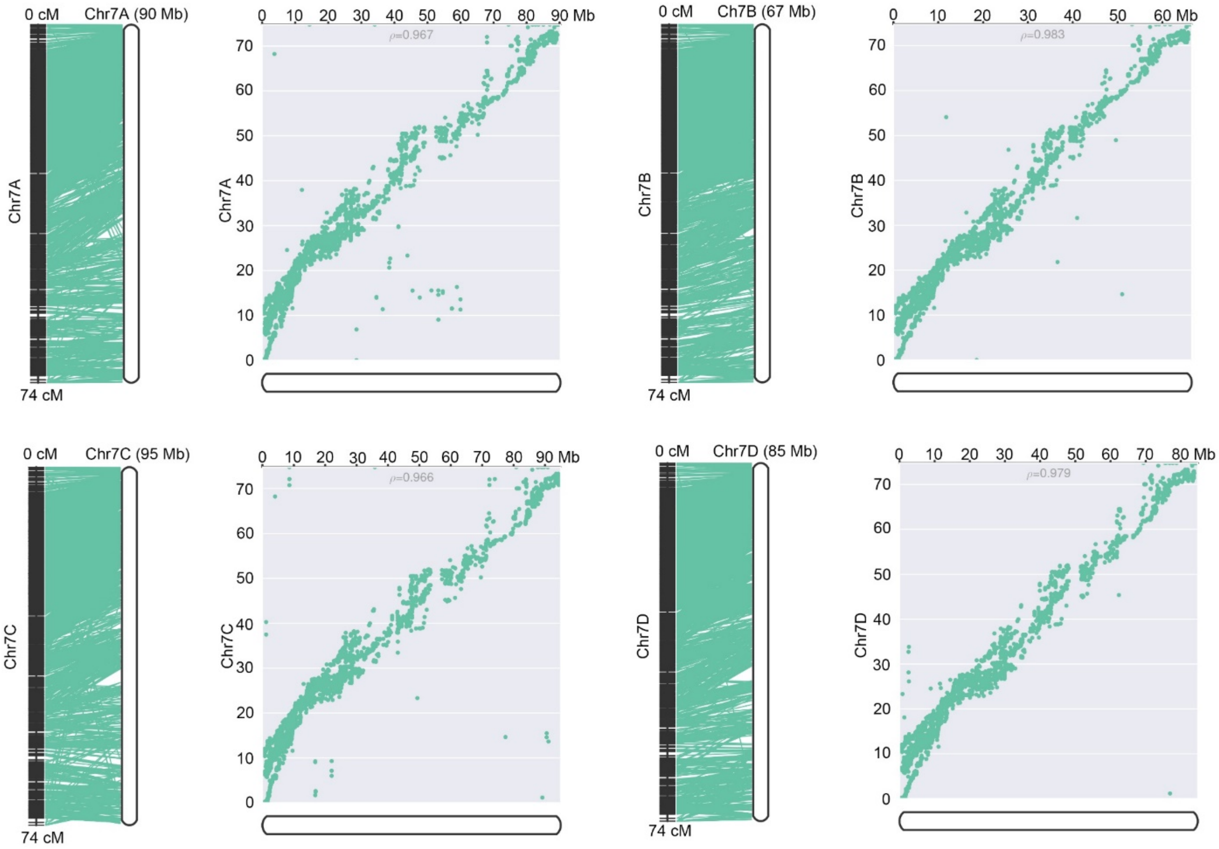


**Supplementary Figure 6.** Continued.

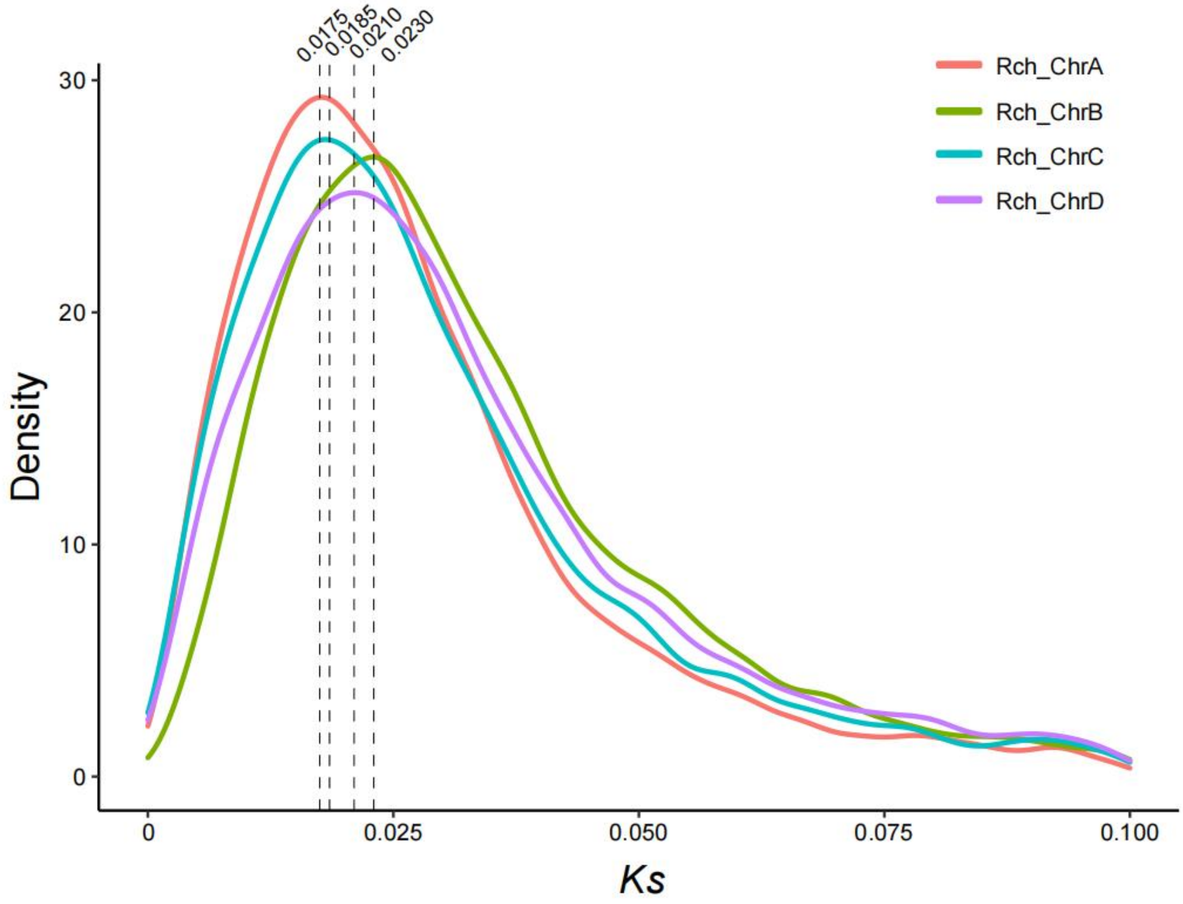


**Supplementary Figure 7.** Distributions of pairwise synonymous substitution rates (*K*s) of orthologous genes between the four haplomes of *R. hybrida* 'Samantha' (ChrA-D) and *R. chinensis* 'Old Blush' (Rch).

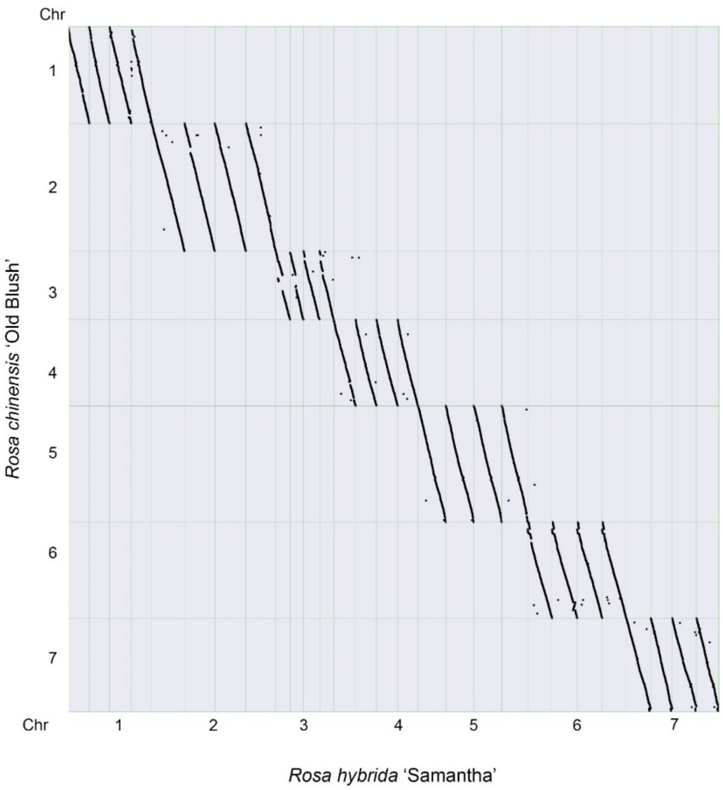


**Supplementary Figure 8.** Synteny plot between the four haplomes of *Rosa hybrida* ‘Samantha’ and the genome of *Rosa chinensis* ‘Old Blush’.

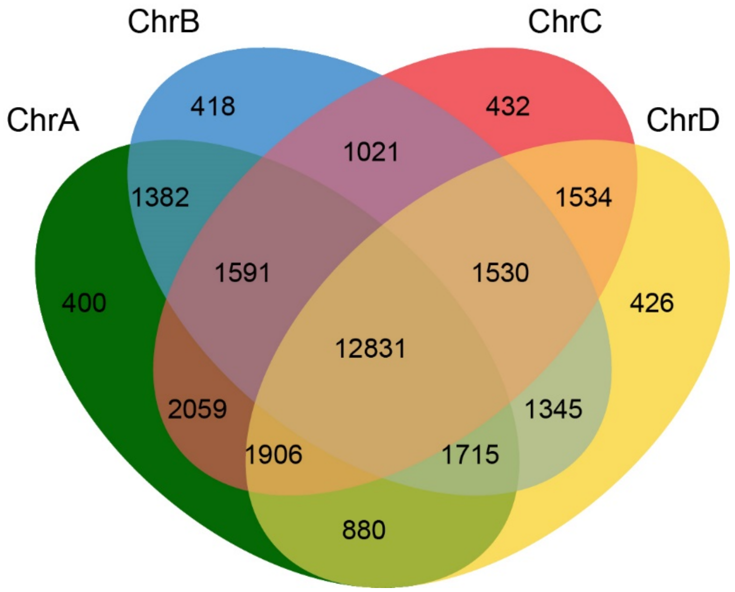


**Supplementary Figure 9.** Gene family clustering analysis of the four haplomes of *Rosa hybrida* 'Samantha'. The Venn diagram displays the intersection of gene families among the four haplomes.

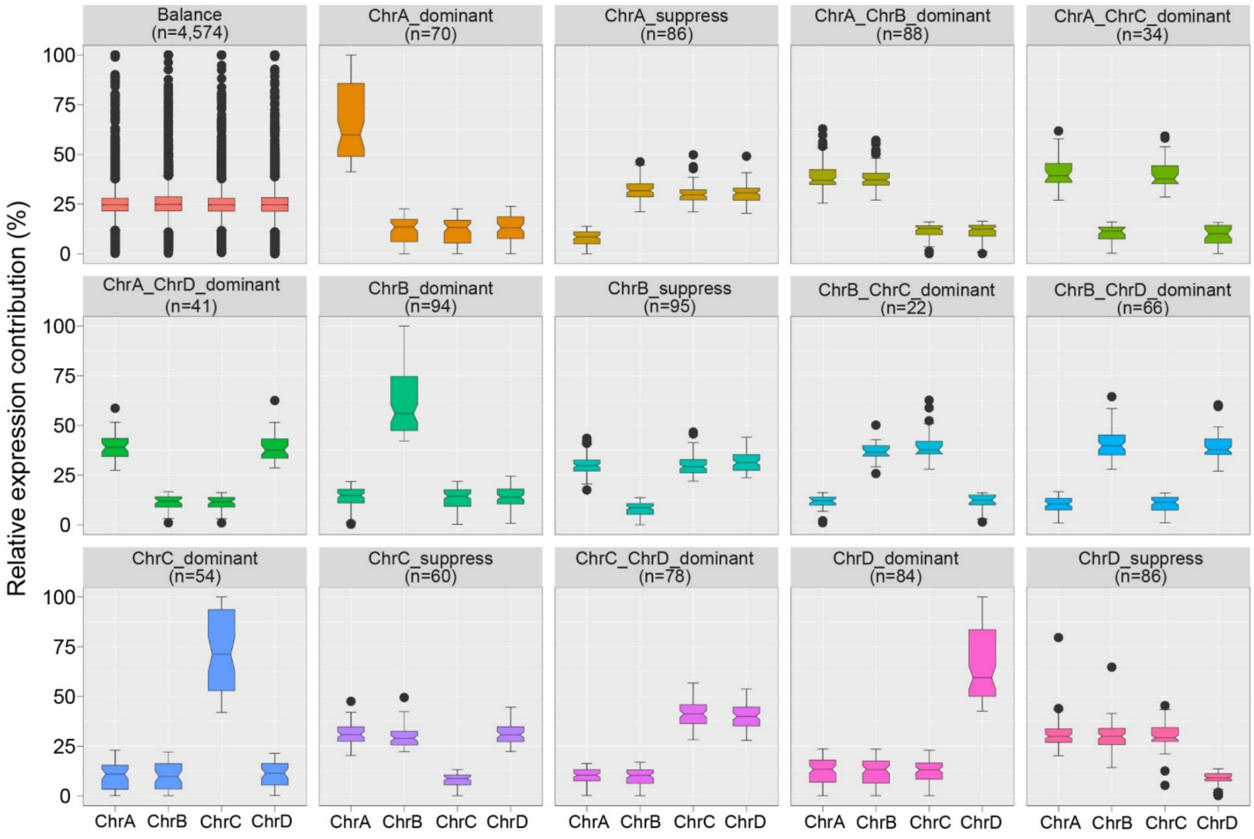


**Supplementary Figure 10.** Expression dominance analysis of syntenic tetrads. The x-axis represents the four haplomes and the y-axis is the relative expression contribution of each syntenic tetrad. ChrA_dominant and ChrA_suppress mean genes on haplome A have higher and lower expression, respectively, than the homologous genes in the other three haplomes; ChrA_ChrB_dominant means genes on haplomes A and B have higher expression than the homologous genes in the other two haplomes; and so on.

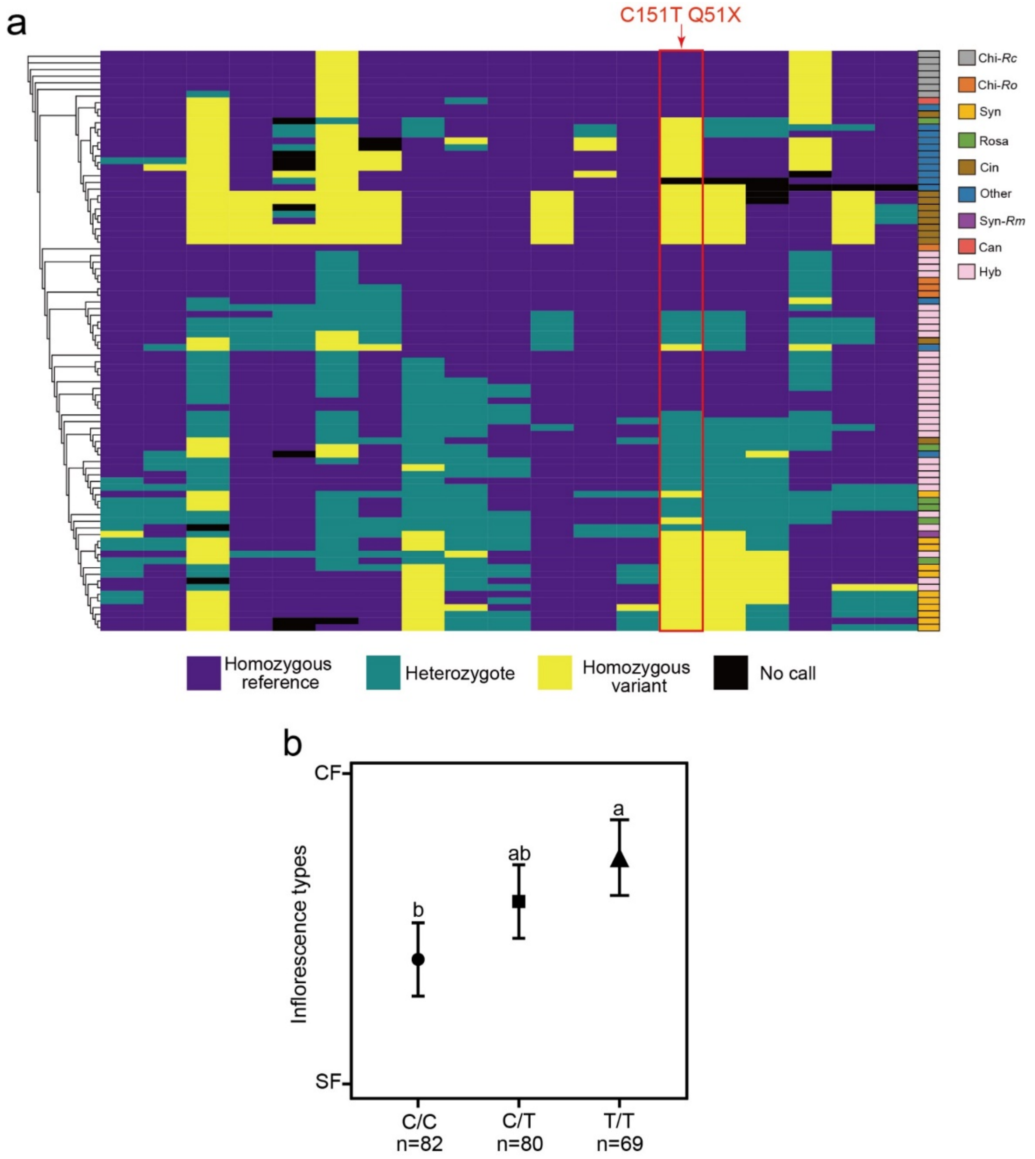


**Supplementary Figure 11. An SNP resulting in a premature stop codon of the *CEN* gene and highly associated with the inflorescence type in modern rose. a,** Genotype analysis of SNPs in the exons of *CEN*. Maximum likelihood (ML) phylogenetic tree constructed using these SNPs is shown on the left. The rectangular shapes on the right indicate the section to which the samples belong. The SNP causing the premature stop is indicated. Hyb, modern cultivars, *R. hybrida*; Syn, section Synstylae; Syn-*Rm*, *R. moschata*; Chi-*Rc*, *R. chinensis* in section Chinenses; Chi-*Ro*, *R. odorata* in section Chinenses; Rosa, section Rosa; Can, section Caninae; Cin, section Cinnamomeae; Other, sections Pimpinellifoliae, Microphyllae, Bracteatae, and Banksianae. **b,** Association analysis between the SNP (Chr6A:53167648) and the inflorescence type in the 233 rose accessions. SF, solitary flower; CF, corymb flower. *P* values were calculated with ANOVA analysis. Different letters indicate statistical significance ($P < 0.05$).

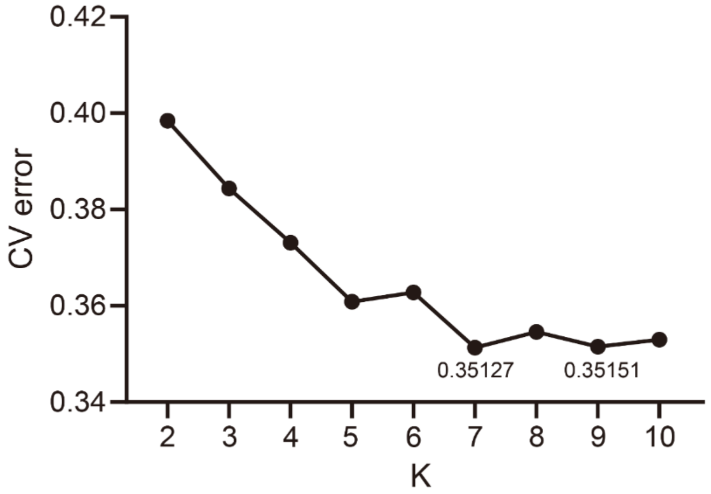


**Supplementary Figure 12.** CV error with *K* ranging from 2 to 10 in the population structure analysis using ADMIXTURE.

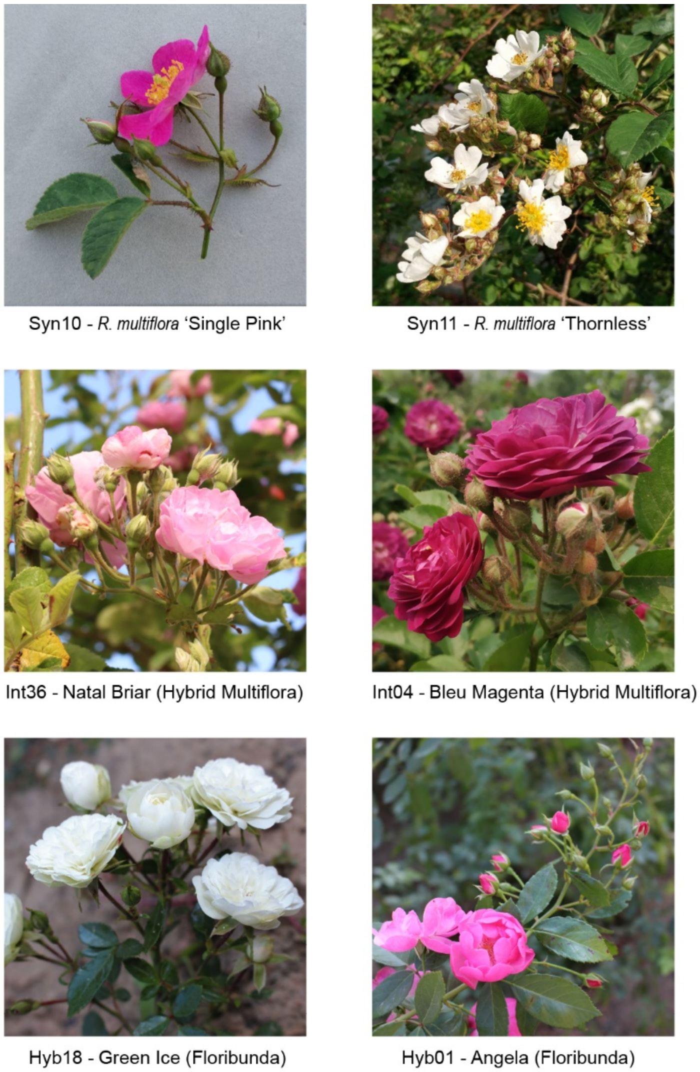


**Supplementary Figure 13.** Inflorescence traits of different rose accessions. The inflorescence trait of the wild species *R. multiflora* has been passed down to Hybrid Multiflora and Floribunda.

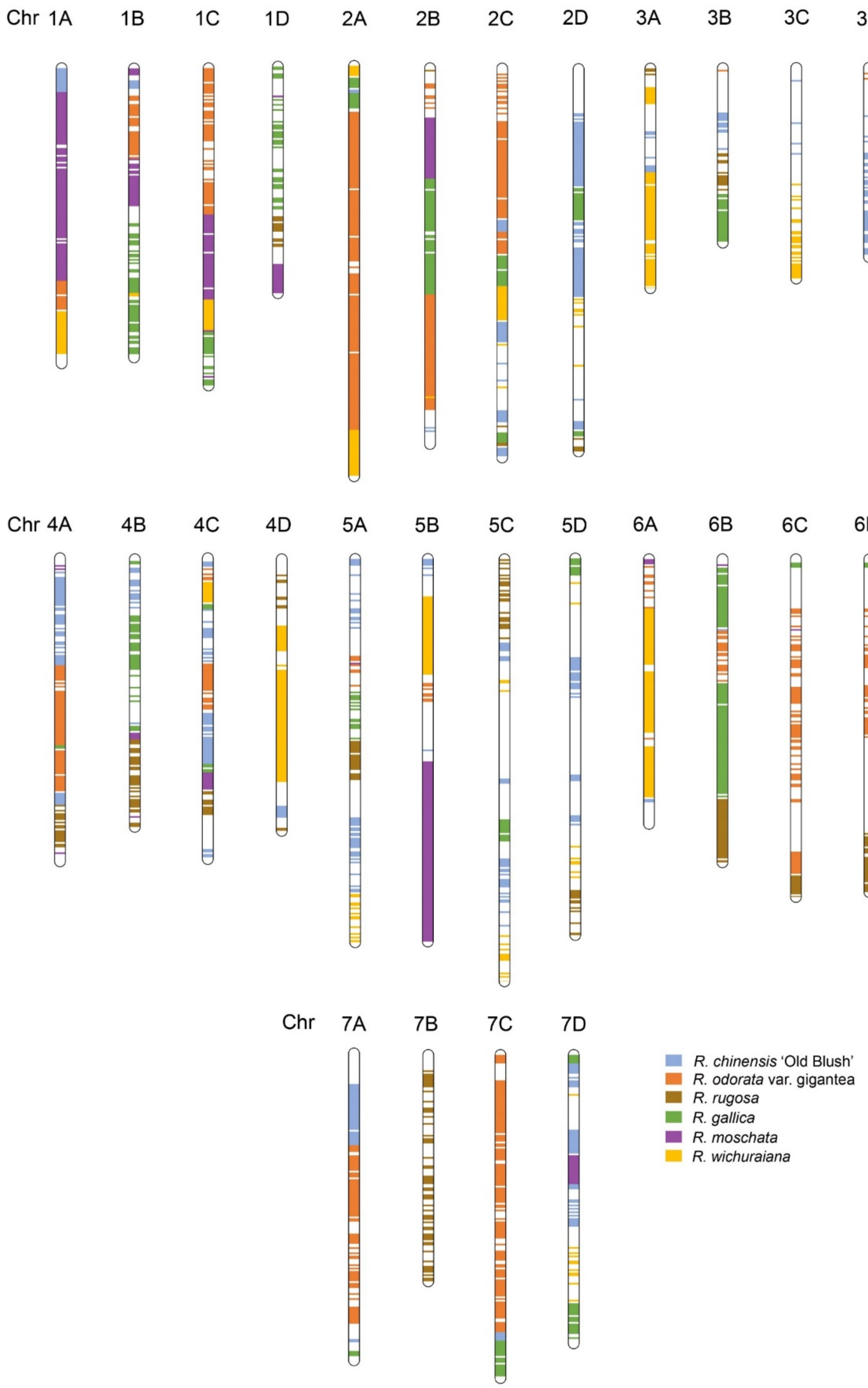


**Supplementary Figure 14.** Genetic organization of the 28 chromosomes of *R. hybrida* 'Samantha'. Different colors represent genome regions derived from different potential original species.

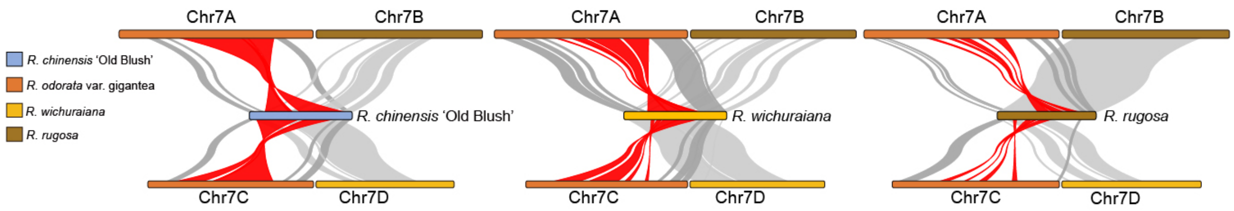


**Supplementary Figure 15.** Collinearity analysis of the region on chromosome 7 harboring an inversion. Three publicly available Rosaceae genomes, *R. chinensis*, *R. wichuraiana* and *R. rugosa*, were used. The potential origins of this region in the 'Samantha' genome are indicated with different colors.

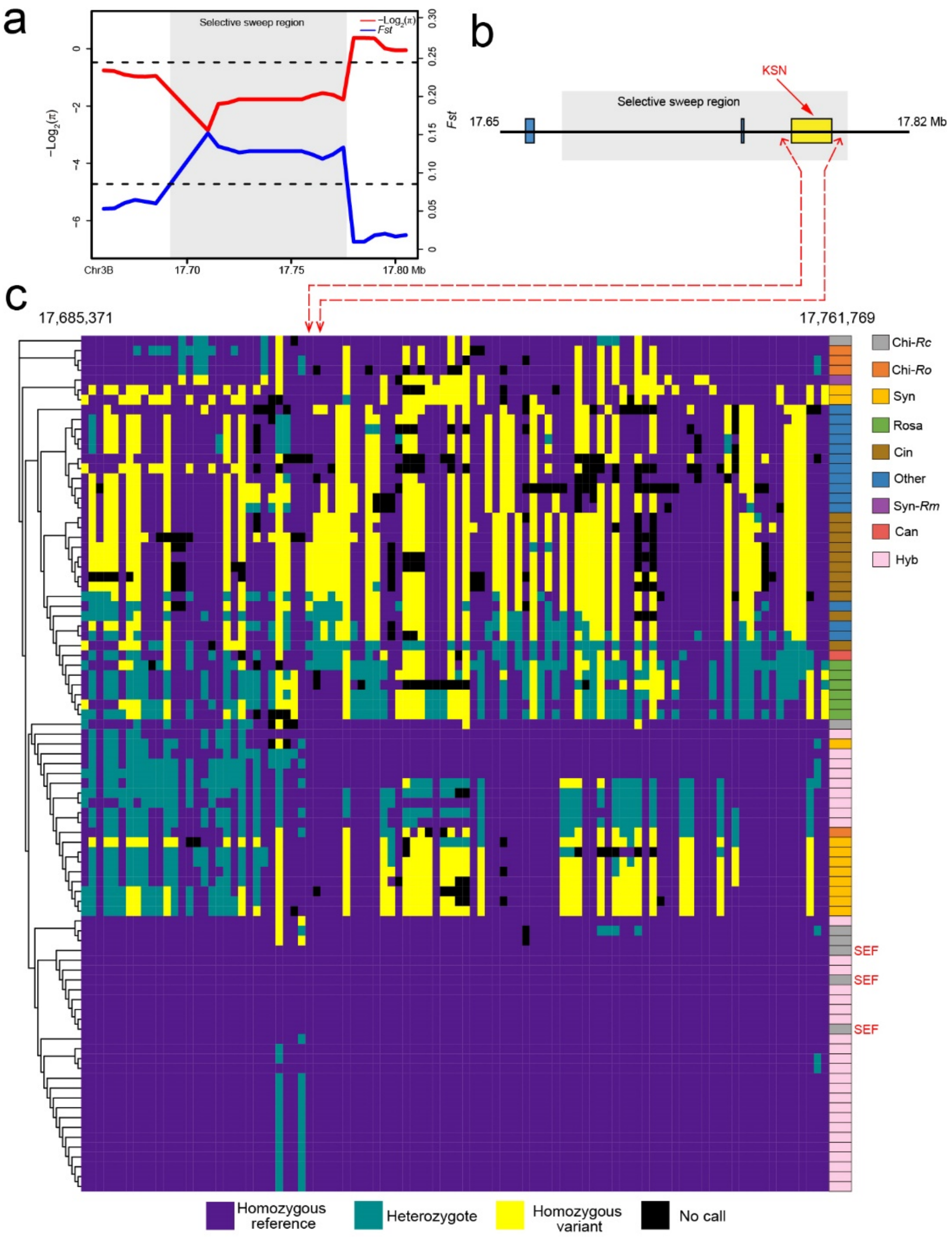


**Supplementary Figure 16. Selection and evolution of recurrent blooming in modern roses.** **a,** π ratio and $F_{ST}$ values in the selective sweep region containing the *KSN* gene. **b,** Genes in the selective sweep region. Blue boxes represent genes located on the sense strand, while yellow boxes represent genes located on the antisense strand. **c,** Heatmap of SNP genotype profiles in the selective sweep region. ML phylogenetic tree constructed from these SNPs is shown on the left. The rectangular boxes on the right indicate the sections to which the samples belong. Hyb, modern cultivars, *R. hybrida*; Syn, section Synstylae; Syn-*Rm*, *R. moschata*; Chi-*Rc*, *R. chinensis* in section Chinenses; Chi-*Ro*, *R. odorata* in section Chinenses; Rosa, section Rosa; Can, section Caninae; Cin, section Cinnamomeae; Other, sections Pimpinellifoliae, Microphyllae, Bracteatae, and Banksianae; SEF, *R. chinensis* 'Semperflorens'.

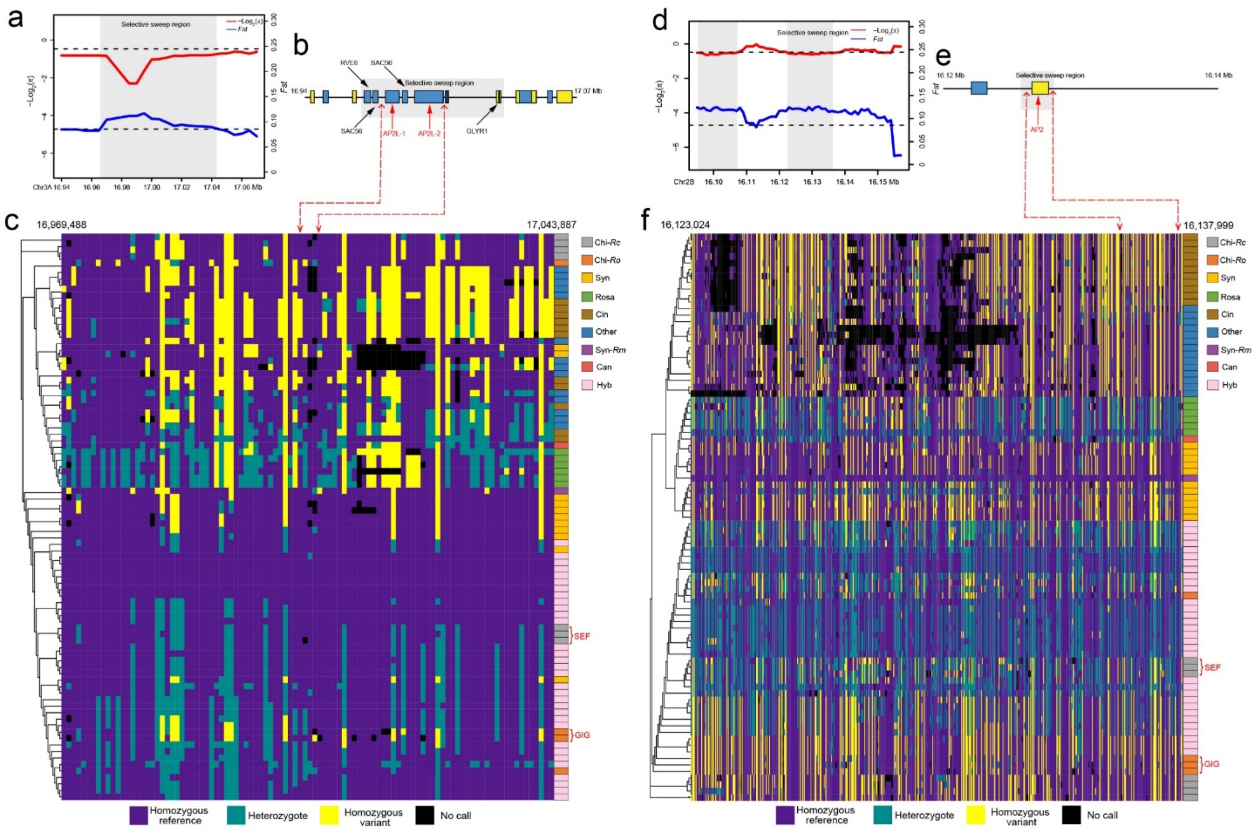


**Supplementary Figure 17. Selection and evolution of double flower in modern roses. a, d,** $\pi$ ratio and $F_{ST}$ values in selective sweep regions containing *AP2L* (**a**) and *AP2* (**d**) genes. **b, e,** Genes in selective sweep regions. Blue boxes represent genes located on the sense strand, while yellow boxes represent genes located on the antisense strand. **c, f,** Heatmaps of SNP genotype profiles in selective sweep regions. ML phylogenetic trees are shown on the left. The rectangular boxes on the right indicate the sections to which the samples belong. Hyb, modern cultivars, *R. hybrida*; Syn, section Synstylae; Syn-*Rm*, *R. moschata*; Chi-*Rc*, *R. chinensis* in section Chinenses; Chi-*Ro*, *R. odorata* in section Chinenses; Rosa, section Rosa; Can, section Caninae; Cin, section Cinnamomeae; Other, sections Pimpinellifoliae, Microphyllae, Bracteatae, and Banksianae; SEF, *R. chinensis* ‘Semperflorens’; GIG, *R. odorata* var. gigantea.

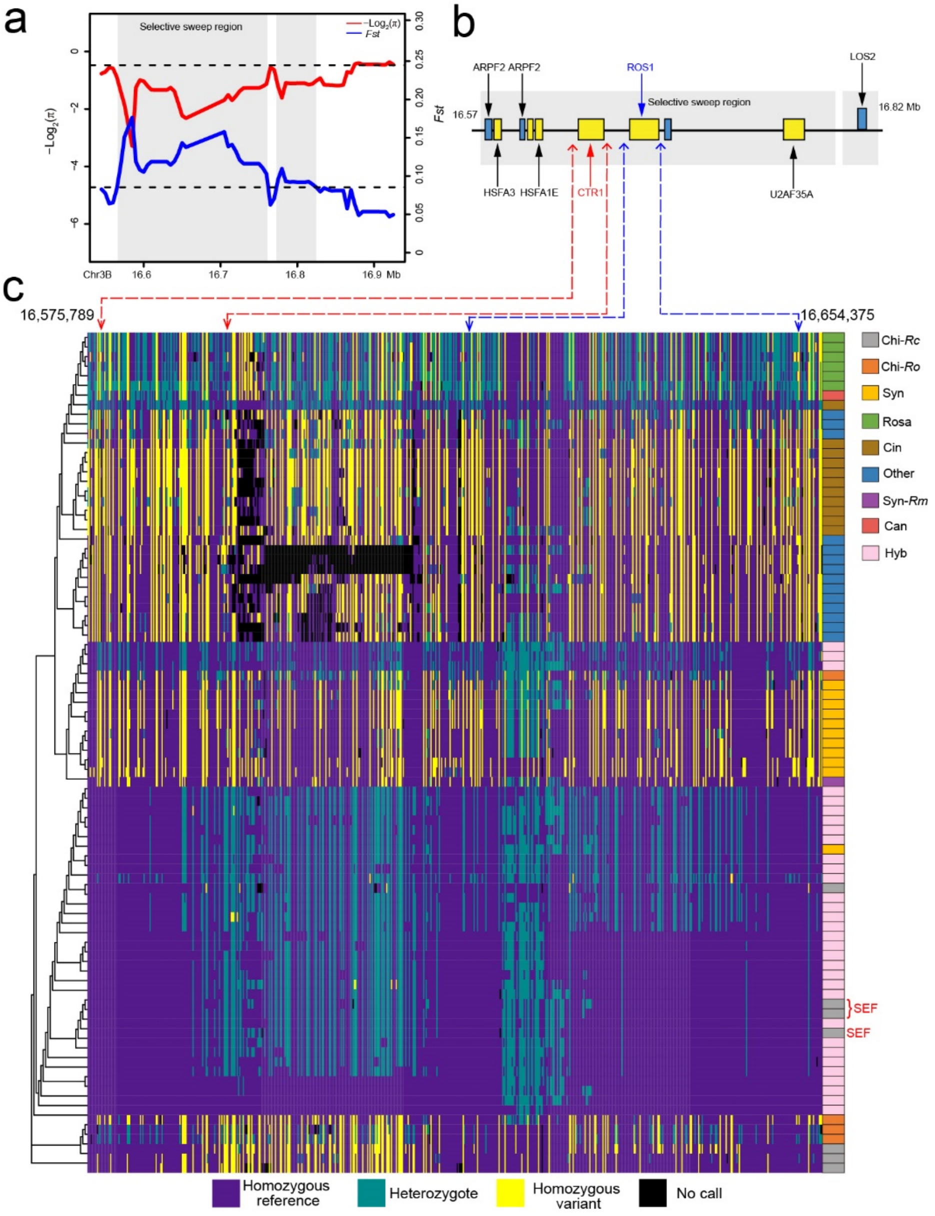


**Supplementary Figure 18. Selection and evolution of ethylene sensitivity in modern roses. a,** π ratio and $F_{ST}$ values in the selective sweep region containing *CTR1 and ROS1* genes. **b,** Genes in the selective sweep region. Blue boxes represent genes located on the sense strand, while yellow boxes represent genes located on the antisense strand. **c,** Heatmap of SNP genotype profiles in selective sweep regions. ML phylogenetic tree is shown on the left. The rectangular boxes on the right indicate the sections to which the samples belong. Hyb, modern cultivars, *R. hybrida*; Syn, section Synstylae; Syn-*Rm*, *R. moschata*; Chi-*Rc*, *R. chinensis* in section Chinenses; Chi-*Ro*, *R. odorata* in section Chinenses; Rosa, section Rosa; Can, section Caninae; Cin, section Cinnamomeae; Other, sections Pimpinellifoliae, Microphyllae, Bracteatae, and Banksianae; SEF, *R. chinensis* 'Semperflorens'.

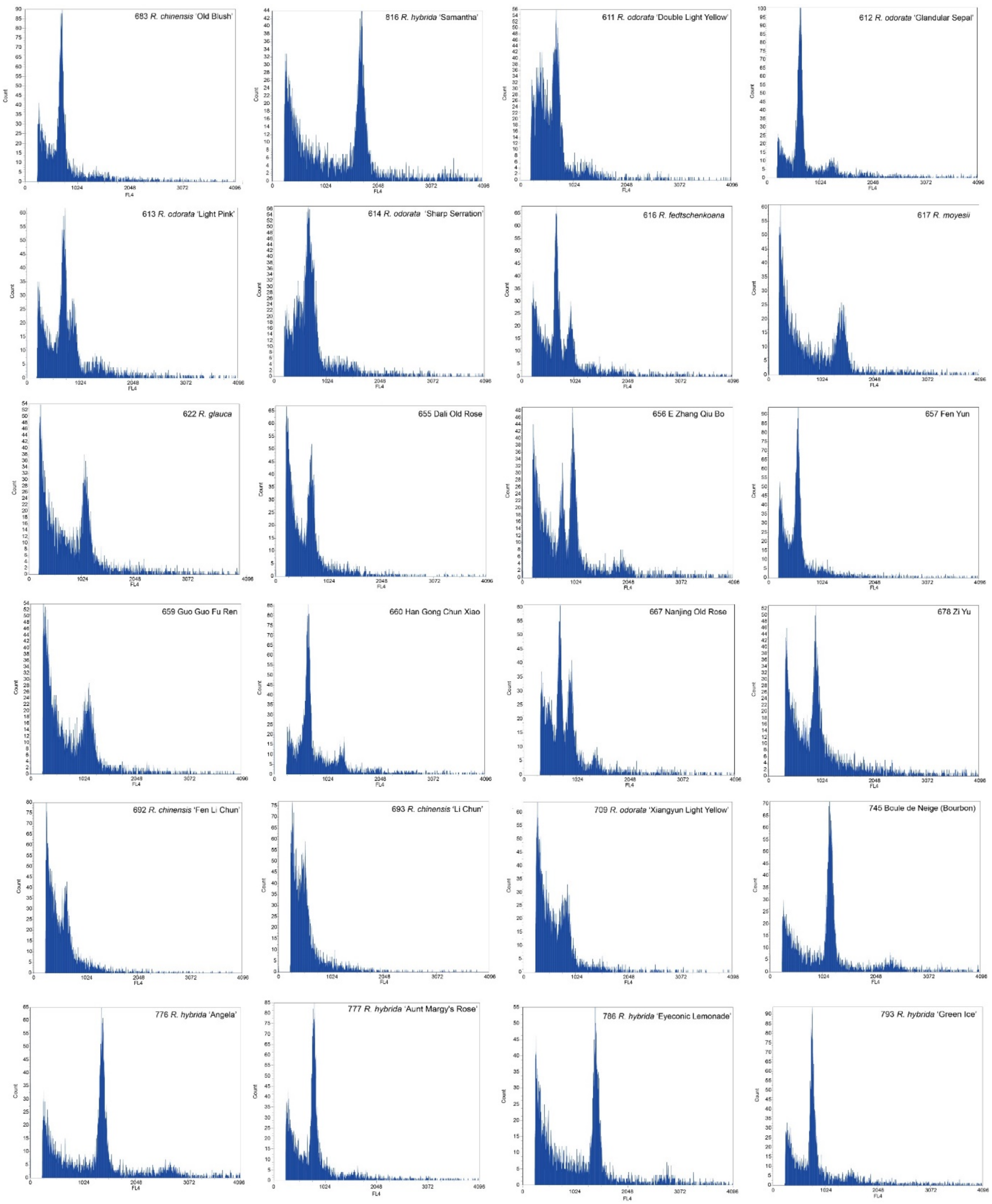


**Supplementary Figure 19. Flow cytometry analysis of ploidy levels in different rose accessions.** The x-axis represents the DNA content, and the y-axis represents cell count. *R. chinensis* 'Old Blush' and *R. hybrida* 'Samantha' are used to represent diploid and tetraploid, respectively.

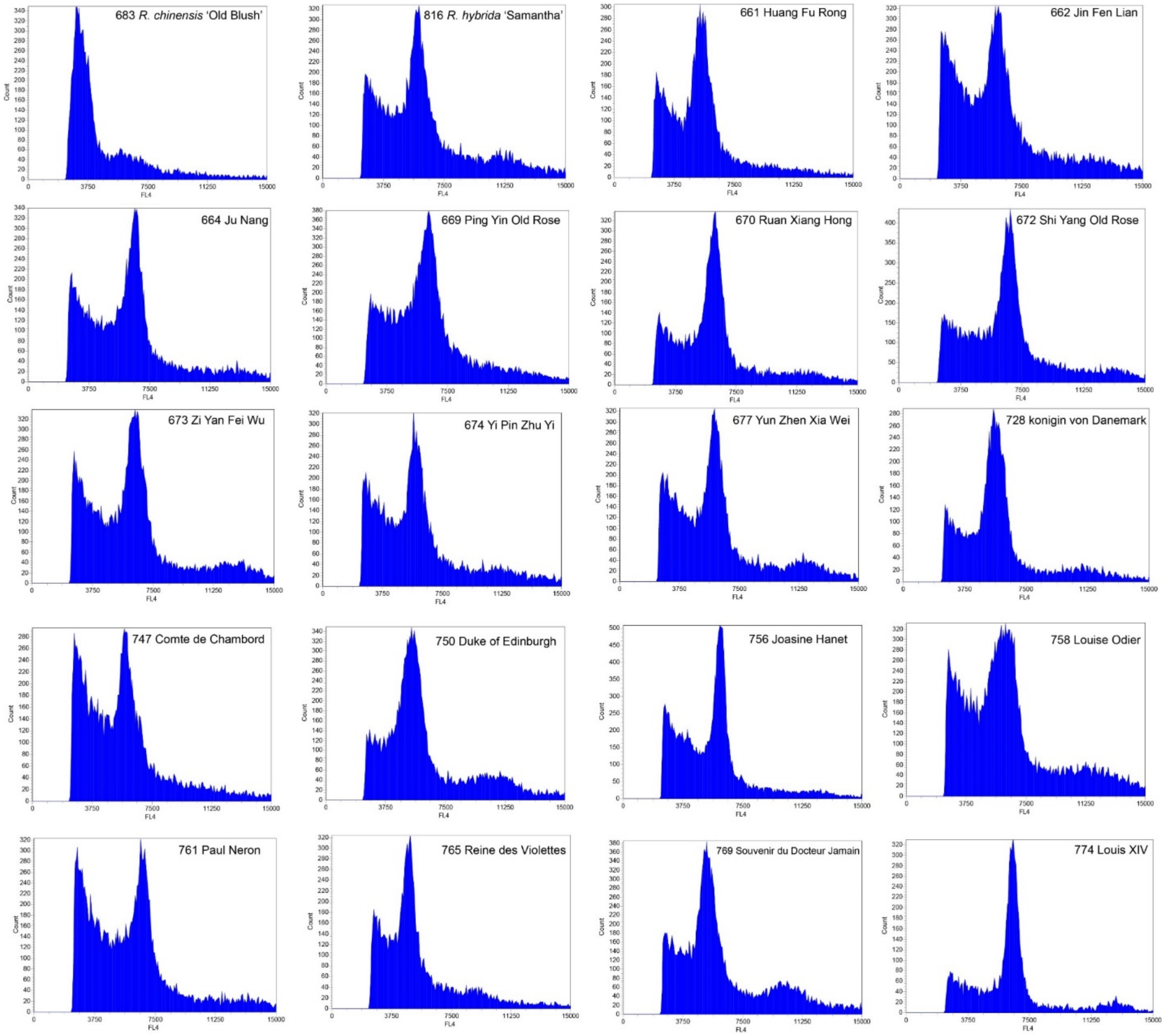


**Supplementary Figure 19.** Continued.

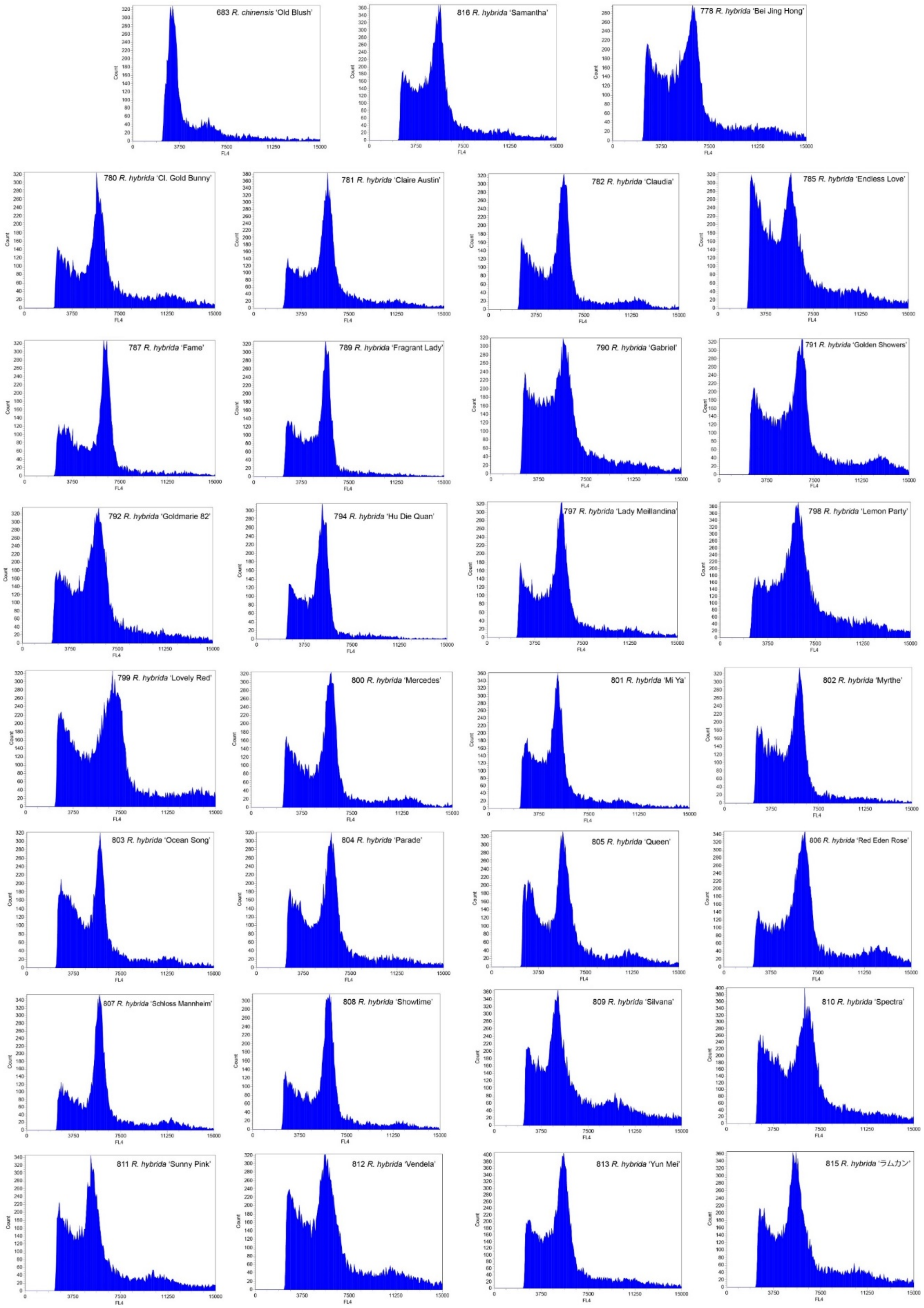


**Supplementary Figure 19.** Continued.
